# Smart utilization of betaine lipids in giant clam *Tridacna crocea*

**DOI:** 10.1101/2023.01.16.524159

**Authors:** Ryuichi Sakai, Naoko Goto-Inoue, Hiroshi Yamashita, Naoya Aimoto, Yuto Kitai, Tadashi Maruyama

## Abstract

The giant clam *Tridacna crocea* inhabits shallow tropical seas with poorly nourished water and severe sun irradiation. They harbor symbiotic algae “zooxanthellae” (dinoflagellate family Symbiodiniaceae) in the mantle tissue and are thought to thrive in this extreme environment by utilizing photosynthetic products from the algae. However, there is no measure of the detailed metabolic flow between the host and symbiont to evaluate one of the most successful symbiotic relationships in nature. Here, we employed liquid chromatography-tandem mass spectrometry (LC-MS/MS)-based lipidomics and Fourier-transform ion cyclotron resonance MS imaging on *T. crocea* tissues, revealing a unique lipid composition and localization with their symbiont algae. We discovered that the non-phosphorous microalgal betaine lipid diacylglycerylcarboxy-hydroxymethylcholine (DGCC) was present in all tissues and organs of *T. crocea* to approximately the same degree as phosphatidylcholine (PC). The fatty acid composition of DGCC was similar to that of PC, which is thought to have physiological roles similar to that of DGCC. MS imaging showed co-localization of these lipids throughout the clam tissues. Glycerylcarboxy-hydroxymethylcholine (GCC), the deacylated derivative of DGCC, was found to be a free form of DGCC in the clams and was isolated and characterized from cultured Symbiodiniaceae strains that were isolated from giant clams. These results strongly suggest that giant clams have evolved to smartly utilize DGCCs, phosphorus-free polar lipids of symbiont algae, as essential membrane components to enable them to thrive in oligotrophic coral reef milieu.

## Introduction

The boring giant clam *Tridacna crocea* is a tridacnid bivalve mollusk and is found in shallow coral reefs. Giant clams are known to engage in symbiosis with the photosynthetic dinoflagellate family Symbiodiniaceae, also known as “zooxanthellae” [1, 2]. The symbiodiniacean cells in giant clams are harbored within a specialized tubular system, “zooxanthella tubes”, which extend from the stomach of the clam and branch out in their mantle region [3]. This symbiotic relationship between giant clams and algal cells has previously been observed from an energy flow viewpoint. For example, algal symbionts within the *T*. *crocea* mantle release glucose, and 46–80% of fixed carbon was translocated from the symbionts to the host tissues [4]. Although these values can vary among studies and clam species, symbiont algae generally contribute more than half of the carbon requirements of giant clams [1, 5]. This implies that giant clams are largely dependent on the symbiont algae for survival, and the symbiotic algae are protected from harmful ultraviolet (UV) radiation by the host clam [6]. Recently, we reported the isolation of ten different natural sunscreen compounds, mycosporines, from the mantle tissue of a giant clam [7]. Mass spectrometry and UV imaging studies have indicated that the distribution of mycosporines within the mantle tissues differs among compounds. This is thought to be related to their UV absorbing function and biosynthetic stage, suggesting that mycosporines can first be biosynthesized by clams or symbiotic algae and then translocated to the area where they can function best with appropriate structural modification [7, 8]. These findings revealed close and complex relationships between the host animal and photosynthetic microalgae, whereby the functional metabolites in the symbiotic system are not just ‘utilized’ as nutrients but are fine-tuned to optimize their functions to thrive in shallow tropical waters. This ‘smart use’ of limited metabolites may have positively contributed to the co-evolution of coral reef invertebrates and their symbiont algae. Symbiotic relationships between invertebrates and microalgae are widely observed in the coral reef environments [9, 10]. Each case of symbiosis is thought to have independently evolved, reflecting their ecological and physiological characteristics. Therefore, many interesting examples of the ‘smart use’ of metabolites are expected in each symbiotic relationship. Although numerous studies on the symbiotic relationship between invertebrate hosts and dinoflagellates have been conducted, these studies have mainly focused on the chemical aspects of carbon and nitrogen flow [11, 12]. Limited studies have investigated the metabolic flow in the symbiotic consortium by identifying individual compounds present in the host and symbiont [13]. Therefore, further understanding of the unique metabolic exchange between the host and symbiont is essential to explain the paradoxical high production in coral reefs. Structural identification of key metabolites by advanced liquid chromatography-mass spectrometry (LC-MS) in combination with the determination of loci by MS imaging may provide a great deal of information regarding the flow and functions of metabolites that govern symbiotic relationships between algae and invertebrates [7]. In the present study, we focused on the lipid component in the giant clam-Symbiodiniaceae system, as lipids can be useful marker molecules for revealing the smart utilization of metabolites, from a metabolic flow perspective, between the host and symbiont. Here, we show the first experimental evidence of the utilization of algal lipids in giant clams. This finding may facilitate further insight into the co-evolution of coral reef organisms on a molecular level.

## Results

### Anatomical analysis of *T. crocea* and distribution of Symbiodiniaceae cells

An anatomical overview of *T. crocea* is shown in Fig. 1. The outermost mantle tissue (epidermal layer; EL) of the clam densely harbors Symbiodiniaceae cells. Conversely, the pale white mantle tissue (inner layer; IL) contains algal cells at a lower density than that observed in the EL. The kidney is a large and dark-colored organ. The adductor muscle and posterior byssal/pedal retractor muscles are located in the middle of the shell. Ctenidia (gill) can be recognized as a comb-like structure. The digestive diverticula (DD) are surrounded by off-white gonads. To avoid cross-contamination as much as possible, each of these parts was carefully separated (Fig. S1). However, DD are embedded in the gonads; thus, a small fraction of the components from interconnected organs was inevitable. To confirm whether algal cells were present or not, we observed algal chlorophyll fluorescence in both the EL and IL of the mantle, muscle, gonad, DD, and kidney in the thin section (Fig. 1 C–H). The algal cells shown as red fluorescent dots under blue excitation light were most densely localized in the EL (Fig. 1C) followed by the IL (Fig. 1D). Although the DD was completely bounded by gonads [14], the DD and gonads could be easily distinguished by color (Fig. S1). The algal cells (including digested cells) were also found in the DD region (Fig. 1H). Although the gonad region was occupied by eggs and sperm (Fig. 1G), small brown patches were observed in rare instances and often contained a few algal cells (Fig. S2). No algal cells were observed in the muscle (Fig. 1F). Although a characteristic granular structure, the nephrolith [15], was observed in the kidneys (Fig. 1E), no algal population was found in this organ. Unfortunately, ctenidia could not be seen in the observed section.

**Fig. 1.**
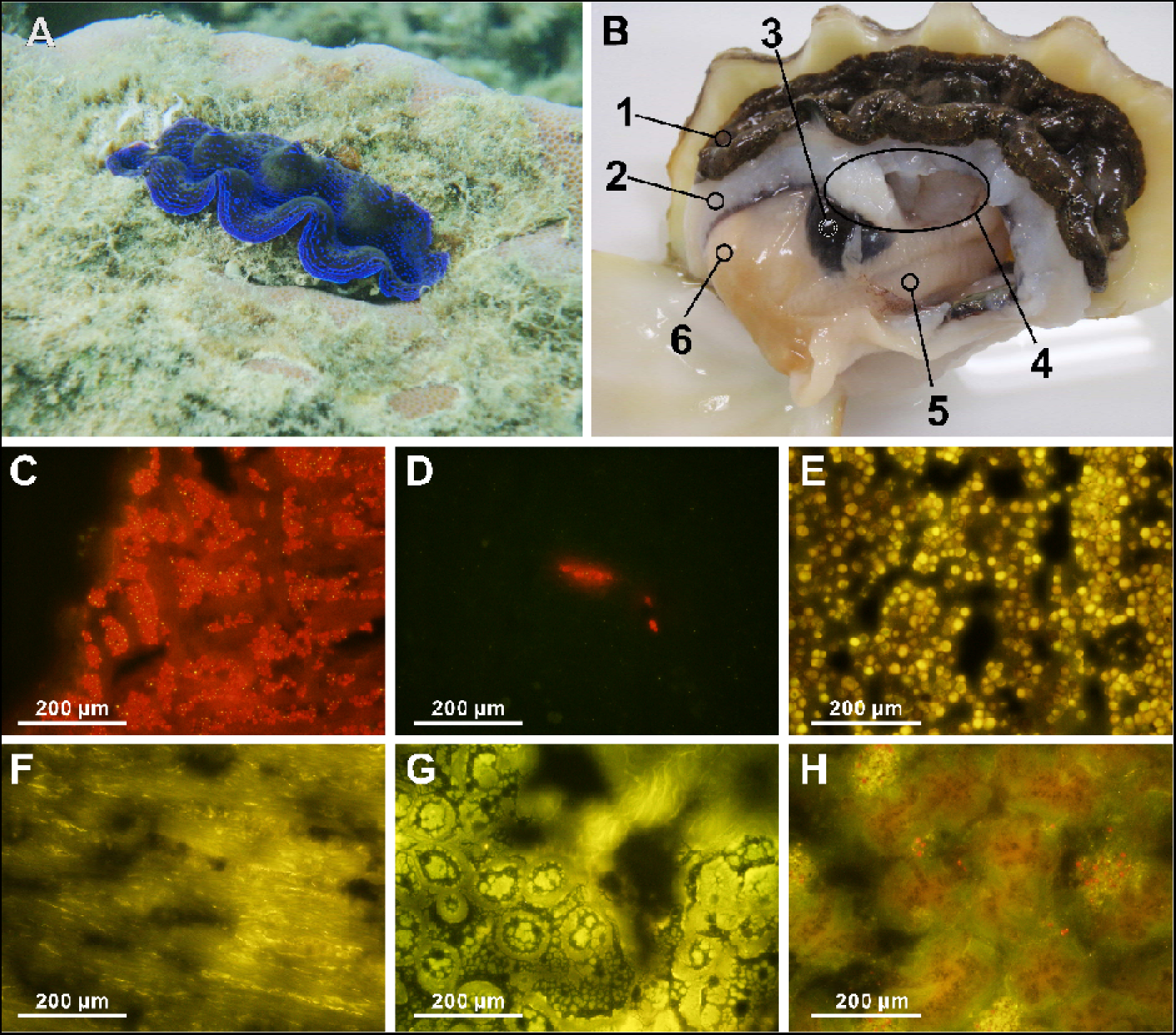
Morphological and histological overview of *Tridacna crocea*. (A) Underwater photograph of *T. crocea* found at – 1 m, at Ishigaki, Okinawa. (B) Dissected specimen (shell length 80.0 mm). 1; mantle (outer epidermal layer: EL), 2; mantle (inner tissue layer: IL), 3; kidney, 4; muscle (adductor muscle and posterior byssal/pedal retractor muscles), 5; ctenidia (gill), 6; gonads and digestive diverticula (DD; inside). (C–H) Fluorescence micrographs of visceral parts of *T. crocea* under blue (460-490 nm) excitation light. (C) Mantle EL, (D) Mantle IL, (E) muscle, (F) kidney, (G) gonad, (H) digestive diverticula. The red dots in fluorescence micrographs (C, D, H) indicate symbiont algal chlorophyll autofluorescence.

### Lipidomic analysis of *T. crocea* organs

We first conducted LC-MS-based lipidomic analyses of extracts from each of the seven organs including mantle (EL and IL), adductor muscle, kidney, DD, gonads, and ctenidia to yield a total of 566 lipids (Table S1). These could be divided into six categories: glycerophospholipids (12 classes, 157 species), betaine lipids (3 classes, 79 species), glycerolipids (9 classes, 209 species), sterol lipids (2 classes, 32 species), sphingolipids (3 classes, 60 species), and free fatty acids (22 species) (Table S1). Semi-quantitative data of the representative lipid classes are summarized in Fig. 2. In all organs, glycerophospholipids were predominant among the membrane-associated lipids (Fig. 2C). Betaine lipids were the second most abundant class and were distributed in all the tissues analyzed, including those that were free of symbiotic algae. The total amounts of glycerophospholipids, betaine lipids, sphingolipids, and sterol lipids were 319, 42, 23, and 11 nmol mg^−1^ dry tissue, respectively (Fig. 2D). To confirm the presence of algal cells in each giant clam body part, we further analyzed algal lipids, namely peridinin and galactosyl lipids, using LC-MS. As expected, peridinin, a dinoflagellate-borne carotenoid was found at a substantial ion intensity in EL and DD, followed by IL (Fig. 2E, Table S2). Only trace ions for peridinin were found in other body parts (Table S2). Moreover, we found three galactosyl lipids: digalactosyldiacylglycerol (DGDG), sulfoquinovosyl diacylglycerols (SQDG), and monogalactosyldiacylglycerol (MGDG) (Table S1, S2). Among them, DGDG was the most abundant class followed by SQDG; MGDG was only detected at low intensities (Fig. 2E, Table S2). Other tissues exhibited negligible peak intensities (Table S2). Subsequently, we semi-quantitatively compared the amounts of phospholipids and betaine lipids at the subclass level in each organ (Fig. 2F). The lipidomic profile of each body part differed substantially (Fig. 2F). Glycerol phosphatidyl ethanolamines (PE) with ether lipids (ether-PE) were predominant among the glycerophospholipid subclasses in all tissues; ctenidia and gonads were rich in this subclass (Fig. 2F). The membrane lipid profiles among muscle, IL, and EL were comparable. Gonads contained considerable amounts of PC and PE in addition to ether-PE. The storage lipid triacylglycerol (TG) was especially prevalent in the gonads (Fig. S3). The kidney was the leanest organ with the lowest phospholipid diversity. Among the three known subclasses of betaine lipids, diacylglycerylcarboxy-hydroxymethylcholine (DGCC) was predominant. Only trace amounts of diacylglyceryltrimethylhomoserine (DGTS)/diacylglycerylhydroxymethylalanine (DGTA) were found in all body parts (10–100 pmol mg^−1^ dry tissue) except for the kidney where less than 10 pmol mg^−1^ was detected. A trace amount of lyso-DGCCs was detected in the ctenidia, DD, EL, gonad, and IL (Table S1).

**Fig. 2.**
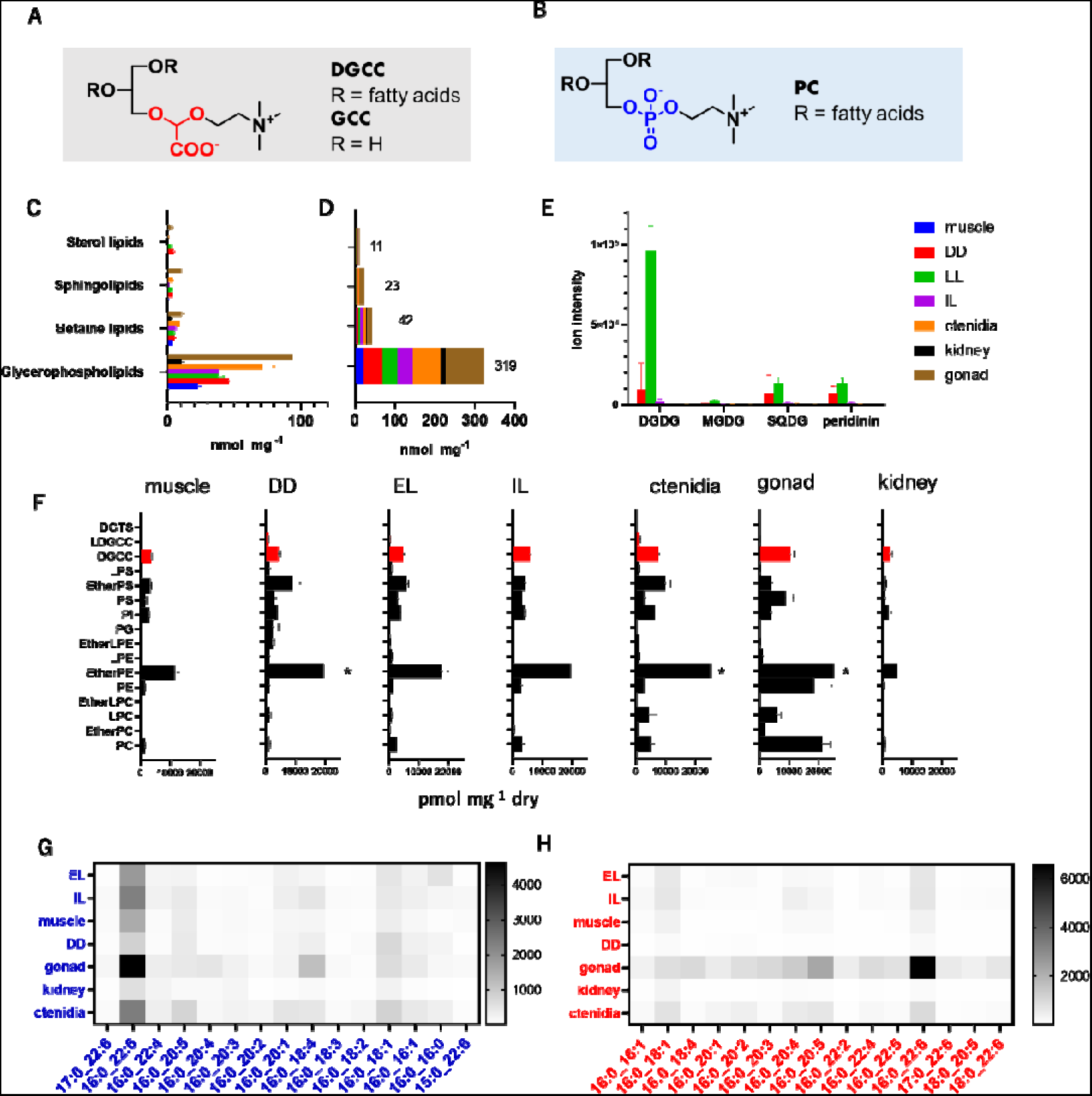
The lipidomics overview of *T. crocea*. Structures of (A) DGCC and (B) PC. (C) Semi-quantitative data for four representative classes of membrane-associated lipids in each organ. (D) Sum of each lipid class. Number on the histogram is the total amount in nmol. Error bars were omitted for clarity. (E) Ion intensity of galactosyl lipids and peridinin in each organ. The color label in the upper light panel is applicable to all graphs in (C–E). (F) Semi-quantitative analysis of the membrane lipids in *T. crocea*. The total amount of each class of lipids in pmol mg^−1^ of dry tissue s shown. Glycerol phosphatidyl lipids, black bars; and betaine lipids, red bars at fixed scale. Some data for ether-PE are clipped* in the given scale (Fig. S3). Comparisons of the 15 most abundant fatty acid species of DGCC (G) and PC (H) in each organ of *T. crocea.* All data here were collected form 5 specimens independently (Table S3, S4). In histograms, average ± SD were indicated.

We next compared the fatty acid composition of DGCC and phosphatidylcholine (PC) because both lipids contain choline as a polar group, and thus they are structurally compatible (Fig. 2A, B). We detected 18 PC and 96 DGCC species, of which the most abundant 15 species were compared (Fig. 2G, H, Table S3, S4). Furthermore, 16:0-22:6 was the major fatty acid in both PC and DGCC. Both classes were comprised mostly of 16:0-containing species (e.g., 16:0-22:5, 16:0-18:1, 16:0-20:5, 16:0-16:1, 16:0-20:1, and 16:0-22:4). Some odd numbered species 17:0-22:6 in both DGCC and PC, 15:0-22:6 in PC were found in *T. crocea* (Fig. 2G, H). Additionally, we analyzed two Symbiodiniaceae culture strains isolated from giant clams (TsIS-H4 and TsIS-G10, Fig. S4), and found that 16:0-22:6 DGCC was the most abundant species in the tested strains as reported previously [16–18]. However, the overall composition of minor species differed greatly from *T. crocea* (Fig. S4). Of note, the PC composition of cultured Symbiodiniaceae cells was substantially different from that of *T. crocea* (Fig. S4).

We also isolated DGCC for further spectral characterization. The ^1^H NMR and mass spectral data for the fraction mainly containing DGCC 22:6_16:0 agreed well with that reported previously from *Pavlova lutheri* [19] (Fig. S5, Fig. S6, Table S5). Moreover, HPLC analysis of DGCC (Fig. S7) supported the data obtained on LC-MS.

### Identification of glycerylcarboxy-hydroxymethylcholine (GCC) in *T. crocea* and cultured Symbiodiniaceae cells

When the aqueous extract of *T. crocea* was analyzed using LC-MS, we found a novel ion at *m*/*z* 252. This ion was absent from the organic extract, suggesting that this compound is a polar water-soluble substance. The molecular formula, C_10_H_21_NO_6_, suggested that the molecule responsible for this ion was the deacylated derivative of DGCC. To obtain pure GCC, we searched for the same ion in cultured Symbiodiniaceae strains; the compound was separated using gel filtration chromatography followed by HPLC. As the proposed structure for GCC has not been previously reported as a free molecule, the planar structure of GCC was determined based on spectral data analyses (Supplementary Results, Scheme S1, Fig. S8-10, Table S5). LC-MS/MS analysis of GCC from Symbiodiniaceae culture strains confirmed the identity between the isolate and that from the clam (Fig. S11).

### Lipidomic analysis of *T. crocea* sperm and fertilized eggs

The symbiodiniacean cells are essential for giant clams; however, fertilized eggs and trochophore stage larvae are free of the symbiont [20], thus the larvae or juveniles must acquire the symbionts from the ambient environment [21]. A subtle decrease in TG ion intensity was observed only after 72 h, while those of DGCC and PC decreased from 24 h and markedly at 72 h after fertilization (Fig. 3A–C). These observations are in line with progress in fertilization as eggs started to divide 3 h after fertilization, while the veliger larvae and D-shaped larvae with thin shells were observed at 24 h and 72 h after fertilization, respectively (Fig. S12).

**Fig. 3.**
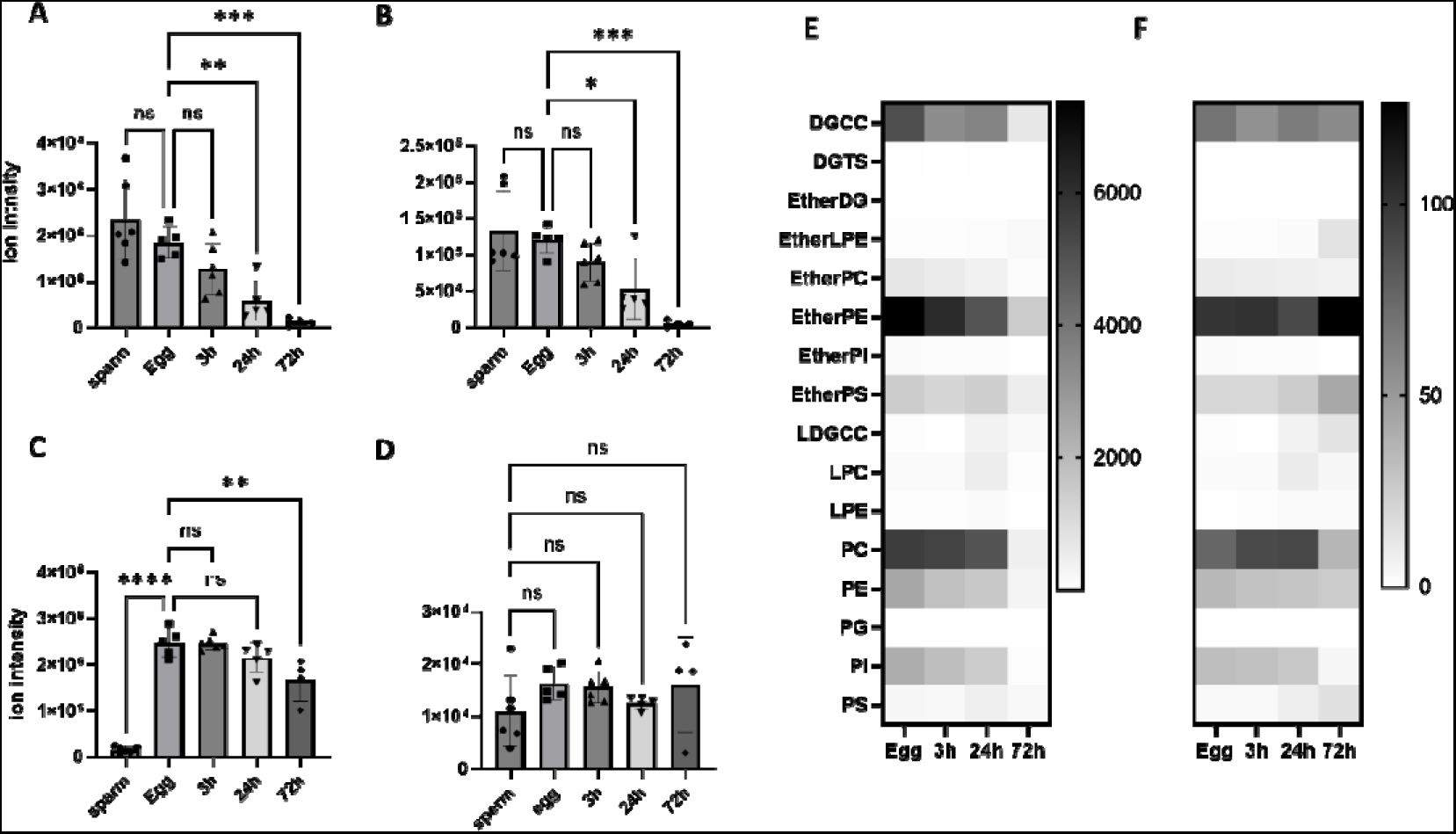
Metabolomics profiles for eggs, sperm, and fertilized larvae of *T. crocea*. Relative abundance of (A) DGCC ** *P* value <0.006, \*\*\*\**P* value < 0.0006, ns: not significant. (B) PC, * *P* value < 0.0215, *** *P* value <0.0004, (C) TG, ** *P* value <0.0011, **** *P* value < 0.0001, and (D) GCC. Each histogram indicates average value from four to six independent specimens ± SD. One-way ANOVA followed by a Dunnett’s test for multiple group comparison, ns: not significant. (E) Semi-quantitative analysis for lipid classes at each developmental stage (pmol mg^−1^). (F) Relative concentration of lipid in each stage. Data in each column were normalized so that the highest value becomes 100 and the lowest value is equal to 0.

Notably, GCC was found at a stable level in all sperm and egg stages (Fig. 3D). Next, we semi-quantitatively compared the membrane lipid composition of eggs and larvae during development. As for the adult clam, the amounts of ether-PE, PC, and DGCC were predominant over other lipids in eggs and larvae up to 24 h. The original compositions of all classes were maintained at 3 h after fertilization (Fig 3E, F). Lipid compositions started to change from 24 h and then changed drastically at 72 h (Fig 3F).

### Lipidomic analysis of other giant clams and bivalves

The above results suggested that DGCC is an indispensable class of fatty acid in *T. crocea*. We thus investigated whether DGCC was present in other giant clams and non-symbiotic clams. To confirm this, we compared DGCC in the adductor muscles of *Tridacna squamosa*, *Tridacna derasa*, and *Hippopus hippopus* along with two other Symbiodiniaceae-free bivalves as controls: *Atactodea striata* and *Donax faba*, which inhabit the sandy shore in Okinawa. The LC-MS data indicated that the giant clams contained DGCC, whereas those in the control group did not have detectable amounts of DGCC (Fig. S13).

### MS imaging analyses

Next, we investigated the localization of lipids in *T. crocea* tissues to assess whether DGCC and PC, structurally compatible lipids, share distribution patterns. The imaging data were analyzed for DGCC and PC with two different fatty acid compositions, 16:0-22:6 and 16:0-18:1, which were predominant molecules in *T. crocea* tissues (Fig. 2). As expected, 16:0-22:6 DGCC and PC showed similar distribution patterns, that is, they were dense in the EL and gonads but only trace levels were visible in the DD, kidney, and muscle (Fig. 4). In the case of 16:0-18:1 species, DGCC was distributed throughout the mantle tissues (Fig. 4B) and visceral parts, including DD (Fig. 4A), while PC showed a similar profile to that of the 16:0-22:6 species, except for some signals that were observed in the IL region of the mantle tissue.

**Fig. 4.**
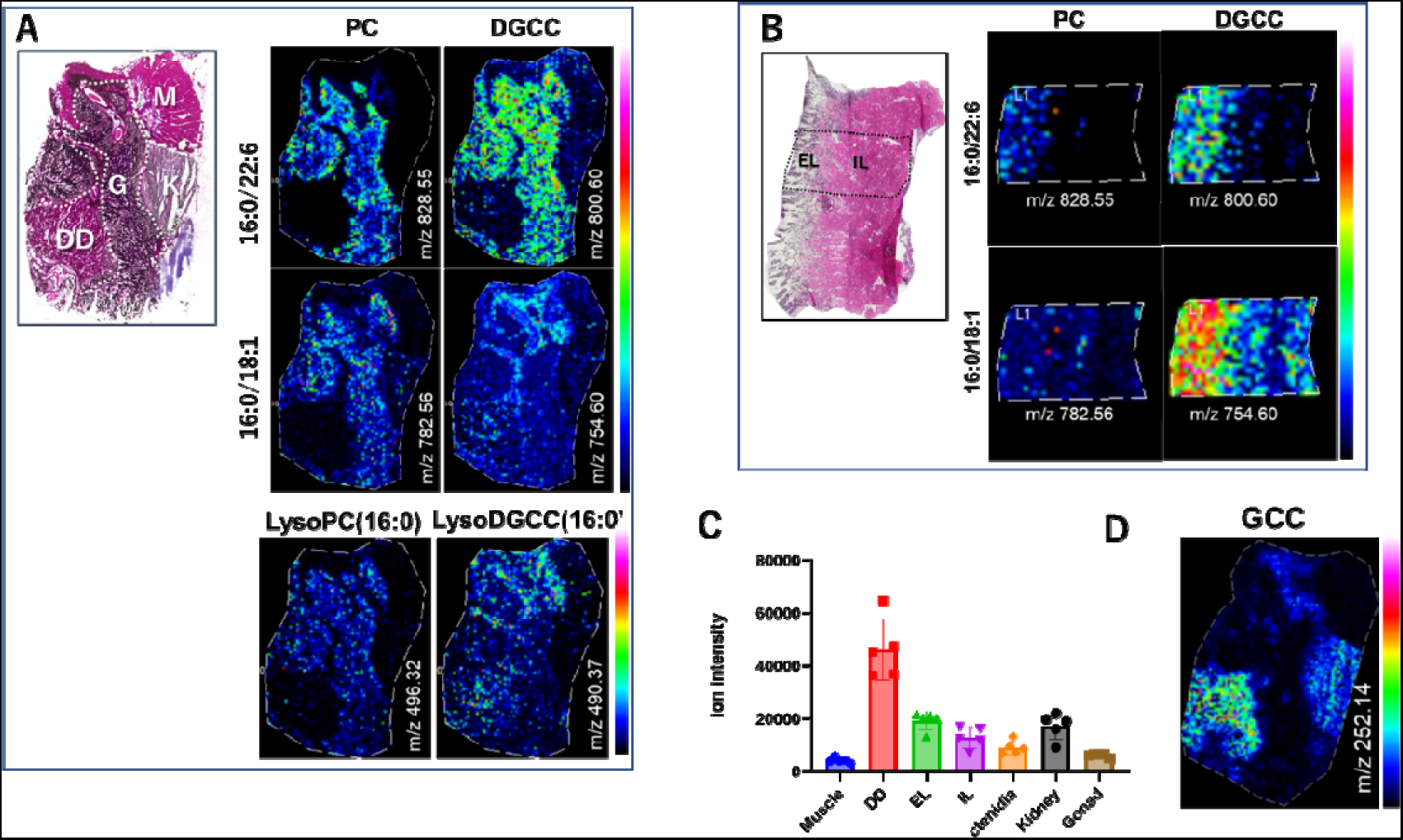
Mass imaging data for cryosections of *T. crocea*. (A) Atlas of the HE-stained cryo-visceral-section recovered after MS imaging. M, muscle; K, kidney; G, gonad; and DD, intestine. Heatmaps (0–100%) show relative intensities for two representative ion species (16:0/22:6) and (16:0/18:1) of PC and DGCC, and lyso-PC (16:0), and lyso-DGCC (16:0) at positive mode, (M + Na)^+^ for PC and (M + H)^+^ for DGCC. (B) HE-stained cryo-mantle-section and heatmaps. (C) LC-MS analysis of GCC in each organ. Each histogram indicates average value form four independent specimens ±SD. Detailed result of multiple comparison is given in Fig. S14. (D) MS-imaging data for detection of GCC.

We then analyzed the distribution of lyso-PC and lyso-DGCC, which can be enzymatically transformed from PC and DGCC, respectively. The distribution pattern of lyso-PC (16:0) was similar to that of PC (16:0-18:1); faint signals were found in the DD, kidney, and muscle. Notably, in DD, signals for lyso-DGCC (16:0) were clearly observed as opposed to those of lyso-PC (16:0) and 16:0-22:6 DGCC; however, the distribution pattern was somewhat similar to that of 16:0-18:1 DGCC.

Further analysis of the distribution of GCC, a deacylated form of DGCC, revealed that its ion distribution was different from that of 16:0-22:6 DGCC, showing dense signals in the DD and kidney, but low intensity in gonads (Fig. 4D). Overall, these observations supported those of the LC-MS analysis of the dissected organs (Fig. 2, Fig.S7).

## Discussion

In the present study, we performed LC-MS lipidomic and metabolomic analyses of the giant clam *T. crocea* and cultured giant clam-associated Symbiodiniaceae to identify the unique metabolic relationships between symbiotic dinoflagellate and their host clams. Strikingly, we found a betaine lipid, DGCC and its polar head group GCC, not only in the algal cells but also in the clam tissue extracts at concentrations comparable to those of PC. To date, three different classes of betaine lipids: DGTS, DGTA, and DGCC have been identified. DGTS and DGTA are structure isomers found in a wide variety of lower plants, such as algae, pteridophyta, bryophyta, lichens, and fungi [22, 23]. However, DGCC is only known in limited taxa of microalgae, including Haptophyceae, Bacillariophyceae, and Dinophyceae [16, 24]. Recently, DGCC was detected in the coral *Montipora capitata* [25] and the zoantharian *Palythoa* sp. [26]; however, the origin of DGCC in these invertebrates is not known since it is difficult to separate symbiont algae from the host to precisely analyze its origin.

We combined metabolomics and MS imaging approaches on large dissectible giant clams to overcome this problem. This combined approach revealed that DGCC was present in considerable amounts in all analyzed tissues of *T*. *crocea*. As expected, DGCC was detected in the Symbiodiniaceae-rich mantle, because the “zooxanthellae” are known to contain DGCC [16]: however, the presence of DGCC in the muscle and kidney, which harbor only a few algae, and in sperm and eggs where no symbiotic algae exist, was unexpected. DGCC is most likely biosynthesized in microalgae [27], and to the best of our knowledge, this is the first report that showed the presence of DGCC in animal cells, except for the above-mentioned Symbiodiniaceae-containing corals [28]. Therefore, the present observations led to a hypothesis that *T. crocea* take up algal DGCC and metabolites in their own cells and tissues. Our data along with previous observations showing that algal DGCC is localized in plasma membranes [29, 30], strongly suggest that DGCC is incorporated in the lipid bilayer of the clam cells, up to a level equivalent to that of PC [23, 31, 32]. The MS imaging data for two major fatty acid species illustrated that the most abundant species 16:0-22:6 of PC and DGCC share similar distribution profiles, suggesting that these lipids are physiologically compatible. However, the second most abundant 16:0-18:1 species of PC and DGCC were distributed somewhat differently (Fig. 4A). This suggests that lipid molecules at the class and even species levels have defined loci to present and roles to play [33, 34]. Phytoplankton in oligotrophic waters with low phosphate concentrations can compensate for this phosphorous requirement by breaking down PC, and concomitantly, fulfill their cellular phosphorus requirements by substituting non-phosphorus membrane lipids for phospholipids [31, 32]. Some marine phytoplankton employ betaine lipids for this purpose [31, 32]. Our results clearly show that *T*. *crocea* accumulate DGCC in all tissues, which strongly suggests that giant clams have evolved to utilize DGCC as a phospholipid substitute to thrive in oligotrophic coral reef waters, similar to phytoplankton. Therefore, the giant clam-Symbiodiniaceae consortium has positively contributed to their co-evolution by compromising atomic scarcity.

The presence of degraded forms of DGCC, lyso-DGCC and GCC, were found in the clams and warrant further discussion. Lyso-DGCC was found in the DD and ctenidia, in substantially higher amounts than in the EL, indicating that lyso-one is probably generated by the enzymatic digestion of DGCC in DD. Notably, significant quantities of lyso-DGCC were found in ctenidia and may suggest that they have specific functions in the loci. One of the notable observations here is the presence and distribution of GCC, which can be both a precursor and catabolite of DGCC. GCC was distributed in the DD and kidney which is a complementary pattern to DGCC (Fig. 4D). This observation supports that the clam digests its own symbiont algae in the DD, and algal DGCC can be degraded to form GCC (Fig. 5). The high GCC signal observed in the DD in the MS imaging data supports this deduction. In fact, degraded algal cells were observed in the DD (Fig. 1H). Fankboner’s detailed microscopic observation of aged algal cells that were culled and digested by amoebocytes and conveyed to DD (35, 36), further supports our finding. Therefore, GCC can be re-acylated to produce DGCC or lyso-DGCC, which is re-distributed and utilized in other organs and tissues as a component of the lipid membrane (Fig. 5). We found that the variation in fatty acid species in DGCC from *T. crocea* was highly diverse (49 species), while the diversity in cultured Symbiodiniaceae strains, which were isolated from giant clams was lower (18 species); however, their profiles differed greatly apart from 16:0-22:6 being the major species in both organisms (Fig. 3F, Fig. S4). This differential fatty acid diversity suggests that the algal DGCC should be re-modeled [37] in the clam to constitute lipids that meet the physiological needs of the animal. Little is known about DGCC metabolism and enzymes involved in the re-modeling of DGCC. However, in *Pavlova lutheri*, cytoplasmic DGCC was shown to act as a carrier of fatty acids to plastid MGDG [29], suggesting the presence of acyltransferase that utilize DGCC as a substrate. Thus, our result suggests that giant clams may have acquired such enzymes and metabolic pathways to utilize algal DGCC. It can be speculated that in addition to DD, the kidney is a key organ for the metabolism of DGCC because a high intensity of GCC was found in the kidney tissue. This indicates that digested GCC was excreted as urine or stored as waste. However, it is thought that the kidney in giant clams associates closely with symbiosis with zooxanthella [15]. Although the function of kidneys in giant clams is not well understood, these organs are disproportionally large, and high enzymatic activity involving proteases is evident [15, 38]. The body plan connecting the DD to other parts such as the gonads, gastrointestinal system, and ctenidia, suggests that the kidneys may play a central role in the use of symbiodiniacean metabolites in giant clams [15, 39].

**Fig. 5.**
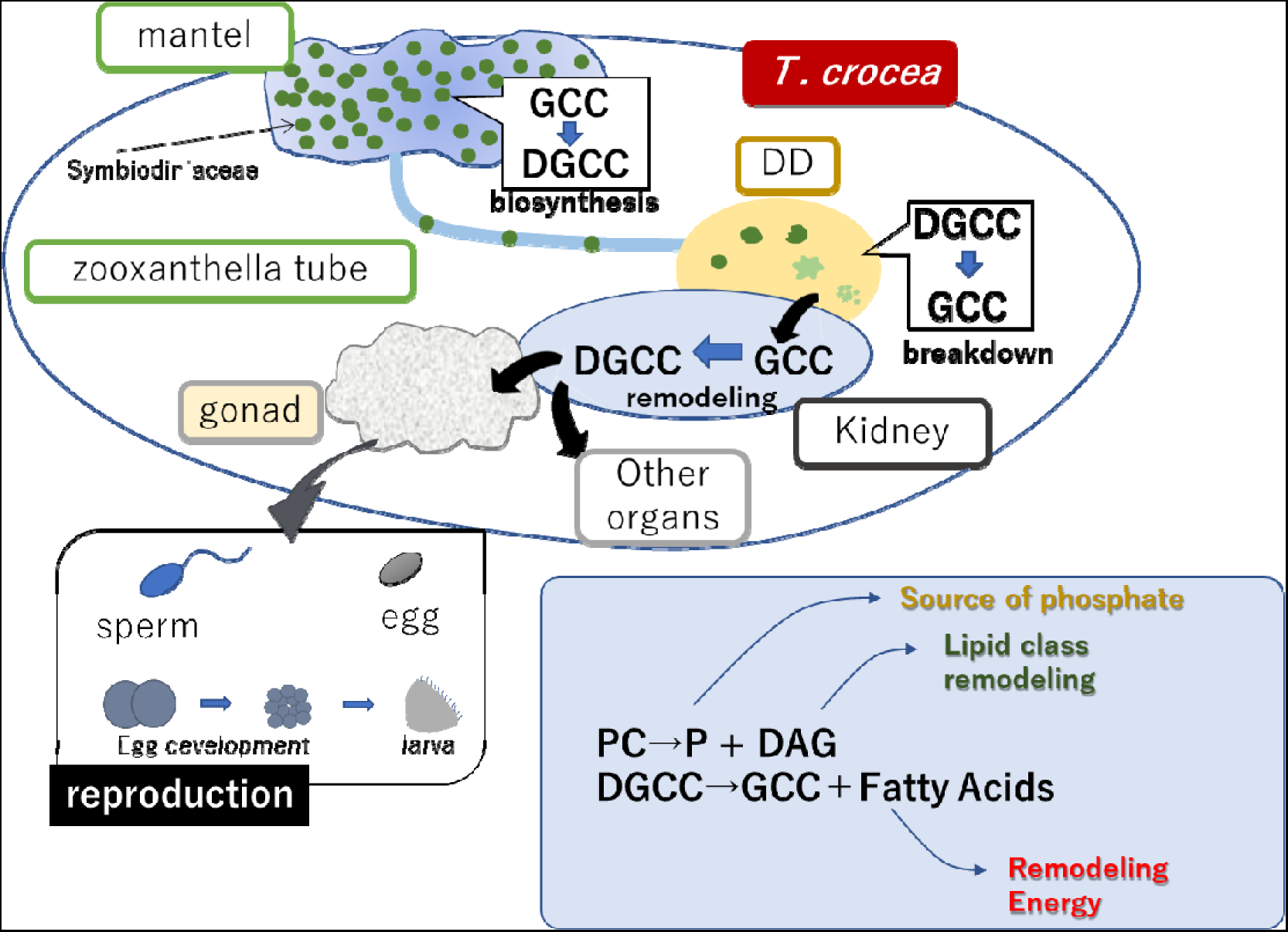
A schematic diagram of metabolic flow in *T. crocea*. Symbiotic algae fix carbons and biosynthesize GCC followed by DGCC. The algal cells are culled, transported, and digested in DD. Algal DGCC can be hydrolyzed to form GCC, which is re-acylated to form DGCC with new acyl substituents (remodeling). Newly formed DGCC are redistributed to each organ to be incorporated in plasm membrane. In phosphorus (P) deficient conditions, the use of DGCC allows consumption of PC as an extra source of P.

The distribution of DGCC in completely algae-free eggs and sperm suggests maternal transfer of these molecules. DGCC was consumed rapidly as development progressed, similarly to PC, while the storage lipid TG was consumed relatively slowly (Fig 3A–C). It was also evident that lipid class remodeling occurred at approximately 24 h after fertilization, suggesting de novo lipid biosynthesis occurs throughout development. The fact that GCC was found in all egg and sperm stages suggests that not only DGCC but also GCC were supplied maternally. The level of GCC in sperm, eggs, and larvae was consistent throughout the stages, suggesting that free GCC can be pooled at certain levels. Although the biosynthesis of DGCC is not understood to date [27, 40], the presence of DGCC and GCC both in Symbiodiniaceae and the clam may provide some clues in investigating the metabolism of DGCC. Taken together, we conclude that DGCC takes part in the membrane lipid metabolism in developing *T. crocea*. It is reported that DGTS take roles of PC in P-deficient conditions and can function as a hub of glycerolipid remodeling in some stramenopile microalgae [41, 42]. Our data suggested that DGCC acts to serve fatty acids in the lipid biosynthesis of *T. crocea*, analogous to other betaine lipids in lower plants.

In conclusion, the lipidomic study combined with MS imaging techniques uncovered an unusual metabolic flow in the giant clam-algae consortium. First, our observation that DGCC was distributed not only within the symbiont-rich sites in the clam but also in the symbiont-free tissues, including eggs and sperm, suggests that the clam uses this lipid as a plasma membrane component. Second, DGCC is most likely enzymatically digested, remodeled, and then re-distributed to organs within the clam. Although additional clam species need to be analyzed in future studies, the non-symbiotic clams tested here did not contain DGCC and GCC, as opposed to all giant clams, which contained DGCC. Therefore, our results support the hypothesis that the DGCC and GCC found in giant clams are closely related to the symbiotic relationship between giant clams and algae. Together, these findings have identified a novel scenario in the metabolic flow between symbiotic algae and clams. First, GCC is biosynthesized within the symbiotic algae and acylated to form DGCC. The algae are trafficked and digested by clams in the DD to generate GCC, which can then be used as a substrate for acyl transferase for remodeling. This transformation may take place in either the DD or kidney. The newly formed DGCC can be re-distributed to other organs as a building block of plasma membranes. Giant clams thus utilize DGCC as a membrane component, which in turn allows the fast breakdown of PC to supply phosphate for other indispensable phosphorus-containing biomolecules (Fig. 5). Our results demonstrated that animal cells can utilize betaine lipids, possibly with unique coupling with phospholipid metabolism, thereby showing that ‘smart utilization’ of novel metabolites helps giant clams thrive in the oligotrophic milieu [32]. These results provide a basis for future investigations into the paradoxical productivity and biodiversity of coral reef ecosystems.

## Material and Methods

### Clam specimens

Specimens of *T*. *crocea* were collected in the coastal waters of Ishigaki Island, Okinawa, Japan under permission from the Okinawa Prefectural Government for research use (No. 30-82, 2-60). *T*. *crocea* specimens were dissected and each organ including the mantle, adductor muscle, kidney, digestive diverticula, and ctenidia (gill) was identified based on a previous report [14]. The mantles were further divided into EL and IL. Other species of giant clams, *T*. *derasa*, *T*. *squamosa*, and *H*. *hippopus* were purchased from local fishers in Okinawa, and Symbiodiniaceae-free clams, *Atactodea striata* and *Donax faba,* were also collected in coastal waters of Ishigaki Island.

### Preparation of sperm, eggs, and larvae

The gonad cutting method was employed to obtain Symbiodiniaceae-free *T*. *crocea* reproductive cells [43]. The gonads were collected from three different *T*. *crocea* individuals. Giant clams are simultaneous hermaphrodites; thus, the gonads consist of both eggs and sperm. Eggs and sperm were separated using a 5 µm mesh filter. Fertilization was conducted using sperm from the other two individuals, under an egg:sperm ratio of 1:50 [44]. After 3 h of fertilization, eggs were washed with 0.2 µm filtered seawater, and were collected. The remaining fertilized eggs were kept in a 27 °C incubator and the water was replaced once a day. The larvae at 24 and 72 h after fertilization were then collected. We repeated this process with another three individuals. Thus, we used a total of 6 sperm, egg, and larvae samples. Each sample was lyophilized and extracted as described below for LC-MS analysis.

### Lipidomics analysis

Total lipid extracts were prepared using the Bligh–Dyer extraction procedure [45]. The lipid layer was analyzed on a High-performance LC-tandem MS (HPLC-MS/MS) using a quadrupole time-of-flight (TOF) mass spectrometer (TripleTOF 5600+; Sciex,) with a BEH C8 column (2.1 × 150, S 2.5 um; Waters). Detailed LC-MS conditions and analysis criteria are described in the SI Materials and Methods. Final data analyses were conducted using GraphPad Prism. Multiple comparisons with one-way ANOVA followed by a Dunnett’s test were used when applicable.

### Isolation of GCC from cultured Symbiodiniaceae cells

Two Symbiodiniaceae culture strains, TsIS-H4, and TsIS-G10, isolated from giant clam *T*. *squamosa* in our laboratory, were used for GCC isolation. The culture strains were maintained in a 27 °C incubator under a light regime of 80–120 μmol photon m^−2^s^−1^ (12:12 h [light: dark] period) in IMK medium (Sanko Jyunyaku, Tokyo, Japan). Each of the cell pellets from these culture strains was extracted with water on ice using an ultrasonic homogenizer. The water extract was separated to give pure GCC, and its structure was determined by spectral analyses as described in the SI Materials and Methods.

### Isolation of DGCC from *T. crocea*

Adductor muscles from eight cultured specimens of *T. crocea* purchased from local fishers were used to obtain DGCC. Details of isolation and structural assignment are described in the SI Materials and Methods.

### MS-imaging

Consecutive 10-μm frozen sections were mounted onto glass slides with/without hematoxylin and eosin (HE) staining and onto indium tin oxide-coated glass slides (Bruker Daltonics, Billerica, MA, USA) for MS imaging. The section unstained with HE was observed under an epifluorescent microscope (BX50, Olympus, Tokyo, Japan). After MS imaging, the sections were subjected to HE staining for morphological observation. Imaging samples were prepared as previously described [46, 47]. MS analysis was performed using SolariX Fourier-transform ion cyclotron resonance (FT-ICR) (Bruker Daltonics) mass spectrometers. Detailed analytical settings are provided in the SI Materials and Methods.

## Statistical Analysis

Data were obtained from randomly collected specimens. Number of specimens is indicated in figure legends. Datapoints representing mean value and error bars ±SD were analyzed using one-way analysis of variance (ANOVA) and Dunnett’s multiple comparison test using GraphPad Prism 8.0.3.

## Abbreviations

DD: digestive diverticula
DGCC: diacylglycerylcarboxy-hydroxymethylcholine
DGDG: digalactosyldiacylglycerol
EL: epidermal layer
IL: inner layer
GCC: glycerylcarboxy-hydroxymethylcholine
LC-MS/MS: liquid chromatography-tandem mass spectrometry
MGDG: monogalactosyldiacylglycerol
PC: phosphatidylcholine
SQDG: sulfoquinovosyl diacylglycerols

## Acknowledgments and funding sources

The authors thank the staff at the Yeyama field station, Fisheries Technology Institute, Japan Fisheries Research and Education Agency for their support during clam sampling. Hiroki Ikeda at Hokkaido University assisted initial data collection. We would like to thank Editage (www.editage.com) for English language editing. This study was supported by Japan Society for the Promotion of Science KAKENHI grant no. 21H04742 to HY.

Some research equipment for the MEXT Project for promoting public utilization of advanced research infrastructure (Program for advanced research equipment platforms, grant number JPMXS0450200120) was used in this study.

## Supporting Information

### Supplementary Methods

#### LC-MS/MS-based lipidomics

Approximately 5 mg of lyophilized samples were homogenized with a mortar and pestle and weighed in 1.5 mL tubes. Then, 200 μL/mg ice-cold extraction solvent (water∶ methanol∶ chloroform 0.8:2:1, v/v/v) containing 1 mg/mL deuterium labeled phosphatidylcholine 15:0–18:1(d7) (PC), lysophosphatidylcholine 18:1(d7) (LPC), phosphatidylethanolamine 15:0–18:1(d7) (PE), lysophosphatidylethanolamine 18:1(d7) (LPE), phosphatidylglycerol 15:0–18:1(d7) (PG), phosphatidylinositol 15:0–18:1(d7) (PI), phosphatidylserine 15:0–18:1(d7) (PS), triacylglycerol 15:0–18:1(d7)-15:0 (TG), diacylglycerol 15:0–18:1(d7) (DG), monoacylglycerol 18:1(d7) (MG), cholesteryl ester 18:1(d7) (CE), sphingomyelin 18:1(d9) (SM), and diacyl-N,N,N-trimethylhomoserine 16:0–16:0(d9) (DGTS) were added to the homogenates and sonicated on ice for 2 min by an ultrasonic homogenizer. The extraction mixture was allowed to stand for 10 min, vortexed, and then centrifuged at 16,000×*g* at 4 °C for 3 min. Next, 200 μL chloroform and distilled water were added to the supernatant (760 µL) and vortexed for 30 s. After centrifugation at 16,000×*g* at 4 °C for 3 min, the upper aqueous layer was removed, and the lower layer was transferred to new 1.5 mL glass vials and evaporated to dryness in a SPEED VAC (Thermo Scientific). The dried extracts were resuspended in MeOH and diluted 10-fold with MeOH for LC-MS analysis.

High-performance LC-tandem MS (HPLC-MS/MS) was performed on a quadrupole time-of-flight (TOF) mass spectrometer (TripleTOF 5600+; Sciex, Framingham, MA, USA) with a BEH C8 column (2.1×150, S 2.5 um; Waters, Milford, MA, USA) and solvents A (0.1% formate + 10 mM ammonium formate in water) and B (0.1% formate + 10 mM ammonium formate in MeOH: 2-propanol (85:15, v/v)). The flow rate was 0.3 mL/min with the following time program: B conc 75%, 0–2 min; 75%–99%, 2–18 min; 99%, 18–24 min; 99%–75%, 24–25 min; 30 min stop. The column oven temperature was 50 °C, and 3 (positive) and 5 μL (negative) of the samples were injected. The following MS parameters were set based on the lipidomic analysis reported previously(1): MS range 100–1,250 Da, ion spray voltage floating +5,5 kV (positive) and −4,5 kV (negative), gas temperature 350 °C, declustering potential 80V, and accumulation time 250 ms. MS2 was measured using the high sensitivity mode for information dependent analysis (IDA), which acquires 15 times MS2 per cycle. MS2 parameters were MS range 100–1,250 Da, gas temperature 350 °C, collision energy 40–70 V (positive) and −30–-60 V (negative). The collision energy in the positive ion mode was stronger than that used by Tsugawa [1] for clearer acquisition of the fatty acid fragment of diacylglycerylcarboxy-hydroxymethylcholine (DGCC).

We used a mixture of all samples for quality control (QC). QC data were acquired for every five samples during the analysis to monitor the intensity drift of peaks detected by LC-MS and reduce the number of failed MS2 acquisitions of peaks in the IDA mode.

#### Lipidomic data analysis

LC-MS/MS data were analyzed using MS-DIAL version 4.24, and lipid annotation was automatically conducted by matching with an *in silico* MS/MS library available on the RIKEN PRIMe website (http://prime.psc.riken.jp/); the results were manually curated with the confirmation of the diagnostic product ion and neutral losses in addition to the fatty acid fragmentation of each lipid species. We used the table reported by Tsugawa to characterize each lipid species [2].

The parameters below were used for the MS-DIAL analysis: retention time begin, 0.5 min; retention time end, 28 min; mass range begin, 100 Da; mass range end, 2000 Da; accurate mass tolerance (MS1), 0.01 Da; MS2 tolerance, 0.025 Da; maximum change number, 2; smoothing level, 3; minimum peak width, 5 scans; minimum peak height 1000; mass slice width, 0.1 Da; sigma window value, 0.5; exclude after precursor ion, true; keep the isotopic ions until 0.5 Da; retention time tolerance for identification, 100 min; MS1 for identification, 0.01 Da; accurate mass tolerance (MS2) for identification, 0.05 Da; identification score cut-off, 70%; using retention time for scoring, true; relative abundance cut-off, 0; top candidate report, true; retention time tolerance for alignment, 0.5 min; MS1 tolerance for alignment, 0.015 Da; peak count filter, 0; remove feature based on blank information, true; sample max/blank average, 5; keep identified metabolites, true; keep removable features and assign the tag, true; and gap filling by compulsion, true.

#### Preparation of sperm and fertilized eggs form *T crocea*

We conducted artificial fertilization using gonad cut method [3] as follows. The gonads were collected from three different individuals of *T*. *crocea*. Eggs and sperm from gonads were separated by using 5 µm mesh filter. Fertilization was conducted using sperm from other two individuals, under eggs: sperm ratio 1:50. After 3 hours fertilization, eggs were washed with 0.2 µm filtered seawater, and we collected the eggs. Remaining fertilized eggs were and kept at 27°C in an incubator changing water once a day. The larvae at 24 and 72 hours after fertilization were also collected. We repeated this process with another three individuals. Thus, we used total 6 sperm, egg, and larvae samples. Each sample was lyophilized and extracted as described above to prepare analyte for LC-MS analysis.

#### Isolation of GCC from cultured Symbiodiniaceae cells

To analyze betaine lipid of Symbiodiniaceae cells, we used three Symbiodiniaceae culture strains (TsIS-H4, and TsIS-G10) isolated from giant clams. The cell pellets from culture strains were extracted with water on ice using an ultrasonic homogenizer (Smurt NR 50M; Microtec, Funabashi, Chiba, Japan). The water extract was separated by dialysis and the small molecular fraction was subjected to Sephadex LH-20 gel filtration chromatography (GE Healthcare, Chicago, IL, USA). The glycerylcarboxy-hydroxymethylcholine (GCC)-containing fraction was further separated using a Hillic HPLC column (Develosil ANIDIUS, NOMURA CHEMICAL CO., LTD, Japan) with an acetonitrile–water gradient, and pure GCC (IH2-78-7, 1.00 mg) was obtained. Subsequently, HRESIMS (*m*/*z* 252.1442 C_10_H_21_NO_6_, D 2 ppm) was performed.

#### Isolation of DGCC of *T. crocea*

Cultured *T. crocea* (8 specimens) were dissected to yield EL (28g), IL (23g), ctenidia (5g), kidney (4g), DD (2g), foot (10g), muscle (6g). The muscle was used to isolate DGCC. Samples were first extracted with 27 mL of extraction solvent system (water:methanol:chloroform = 2 : 5 : 2) and water (12mL) was added after homogenization. The solvent was partitioned by centrifugation and both upper layer and lower layer were concentrated to dryness. The organic extract (lower layer, 11 mg) was separated by counter current partition chromatography (CPC MODEL LLB; SIC Japan, Tokyo, Japan) using a solvent system; chloroform: n-heptane: n-butanol: methanol: acetic acid (60%) at 3:5:3:3:5 ratio. The lower phase was used as the mobile phase at a rotor speed of 1000 rpm. The sample was suspended in the upper and lower layers of the solvent system (3 mL) and injected in the descending mode at a flow of 9.0 mL/min. Elution was initially performed in the descending mode at 2 mL/min with a column pressure of 35 kg/cm² and then switched to the ascending mode (9 mL/min, 22 kg/cm²). The column was then flushed with methanol, after which the eluents (31 tubes, 10 mL/tube) was collected and combined into 10 fractions (IH9-28-1∼10). Each fraction was analyzed by thin layer chromatography and LC-MS/MS to characterize its lipid composition. Fraction 3 (IH9-28-3, 4 mg) containing DGCC 22:6_16:0 (HRESIMS *m*/*z* 800.5954, M + H^+^, C_10_H_21_NO_6_) as a major lipid was used for spectral analysis.

#### Analysis of DGCC on HPLC-CAD

As DGCC does not offer strong and characteristic UV absorption, charge aerosol detector (CAD, CORONA Ultra RS; Thermo Fisher Scientific, Waltham, MA, USA) was used to quantify DGCC to further support our LC/MS analysis data. We used the lipophilic portion of the dissected samples described above. Each of the extract was dissolved in MeOH (1mg/mL), and POPG was added as an internal standard at final concentration of 1mg/mL. InertSustain C8-5 (4.6 x 250 mm, GL Sciences) was used with a gradient of acetonitrile (0.05% TFA) and water (0.05% TFA) at oven temperature at 50°C. Amount of DGCC was calculated relative to that of POPG (Fig S5).

#### MS imaging

Consecutive 10-μm sections were cut directly from the frozen samples using a cryostat (CM 1950; Leica Microsystems, Wetzlar, Germany). Serial sections were mounted onto glass slides for with/without hematoxylin and eosin (HE) staining and onto indium tin oxide-coated glass slides (Bruker Daltonics, Billerica, MA, USA) for MS imaging. The section without HE stain was observed under epifluorescent microscope (BX50, Olympus, Tokyo, Japan) to confirm the distribution/localization of Symbiodiniaceae cells in the section. After MS imaging, the sections were subjected to HE staining for morphological observation. Samples were prepared as previously described [4, 5]. Briefly, a matrix solution containing 50 mg/mL 2,5-dihydroxybenzoic acid in methanol: water (8:2, v/v) was used, with 1–2 mL prepared before use and sprayed uniformly over the frozen sections using an airbrush with a 0.2-mm nozzle (Procon Boy FWA Platinum; Mr. Hobby, Tokyo, Japan). MS analysis was performed using TOF/TOF 5800 (AB Sciex, Framingham, MA, USA) and SolariX Fourier-transform ion cyclotron resonance (FT-ICR) (Bruker Daltonics) mass spectrometers. To optimize FT-ICR MS, we set the mass range from *m*/*z* 400–1200 for DGCC/PC, and *m*/*z* 100-500 for GCC and the spatial resolution to 150 μm for mantle tissue and 220 μm for the frozen viscera of the animal.

### Supplementary Results

Structure determination of GCC and identification of DGCC.

The molecular formula for GCC, C_10_H_21_NO_6_ was established by the HRESIMS molecular ion at *m*/*z* 252.1442 (M + H)^+^, D 2 ppm. ^1^H NMR (Fig. S6, Table S5) data indicated a presence of trimethyl amino group, and a characteristic singlet appeared at d4.85. In the ^13^C NMR data only 6 resonances (Fig. S7), accounting for 8 carbons appeared. Resonances for C-1” and carboxylate, were missing. Two-dimensional NMR data, however assigned most ^1^H and ^13^C resonances as shown in Table S5. HSQC spectrum was particularly useful to detect acetal carbon 1” which was missing in ^13^C NMR data. Thus, these data constructed the planer structure of GCC mostly, except for a carboxylate group. ESIMS/MS data showing diagnostic ions for DGCC at *m*/*z* 104 and 132 (Scheme S1, Fig. S8, 9), however, supported the proposed structure with the carboxylate group at C-1”. Thus, the planer structure of GCC was assigned.

The molecular formula for DGCC 22:6_16:0, C_10_H_21_NO_6_ was supported by HRESIMS *m*/*z* 800.5954, M + H^+^, D7 ppm (Fig. S10). ^1^H NMR data showed characteristic acetal proton at d4.78 (Fig. S11), however, area of this peak and trimethylammonium group was about 1:24, suggesting about the half of this fraction contains PC. ESIMS showing diagnostic fragment ions at *m*/*z* 104 and 132, assuring the identity of this lipid.

### Supplementary Scheme

**Scheme S1. Planer structure of GCC.** (A) fragmentations in ESIMS/MS. (B) Correlation NMR data for GCC.

### Supplementary Figures

**Fig. S1**. Dissected clam (*Tridacna crocea)* specimen, indicating all body parts used for LC-MS analyses. Adductor muscle is covered by a thin reddish-brown membrane containing Symbiodiniaceae cells. To avoid contamination of algal cells, we removed this membrane from adductor muscle specimens.

**Fig. S2. Light micrographs for gonad region.** Small brown patches (arrows) were observed in the gonad region. B, C high magnification observations. Under blue right excitation, a few algal cells were recognized within the patches (C).

**Fig. S3. Amounts ether-PE (left) and storage lipids TG (right).**

**Fig. S4. Heatmap for DGCC (left) and PC (right) in Symbiodiniaceae culture strains (TsIS-H4, and TsIS-G10).** Positive ion mode was used for DGCC analysis while negative mode was employed for PC data acquisition because substantially fewer peaks were obtained in the opposite polarity in both classes.

**Fig. S5. ^1^H NMR (400 MHz) for the DGCC-containing fraction taken in CD_3_OD**

**Fig. S6. ESIMS for DGCC 22:6_16:0**

**Fig. S7. CAD analysis of DGCC in each body part of *T. crocea* (A) HPLC-CAD trace of each extract. A large peak at *T*_R_ around 22 min is POPG (1 mg/mL), and DGCC eluted right after the standard (black arrow). The area of concentration of DGCC was calculated from the area of standard and DGCC. One sample was employed to obtain this data due to sample availability.**

**Fig. S8. ^1^H NMR (400 MHz) of GCC**

**Fig. S9. ^13^C NMR (100 MHz) of GCC in D_2_O**

**Fig. S10. ESIMS/MS data for GCC**

**Fig.S11 LC-MS analyses of GCC from *Symbiodinium* (A) and *T. crocea* (B)**

**Fig. S12. Eggs, sperms, and larvae of *Tridacna crocea* in each developmental stage**

**Fig. S13. Ion intensities for DGCC in giant clams and other bivalves. *T. squamosa* (n = 3), *T. derasa* (n = 3), *Hippopus hippopus* (n = 1), and bivalves *Donax faba* (n = 3), and *Atactodea striata* (n = 3).**

**Figure S14. Ion intensity for GCC in each organ.**

N = 5, One-way ANOVA followed by a Dunnett’s test for multiple group comparison. *P* values are indicated.

### Supplementary Tables

**Table S1. A summary of semi-quantitative lipidomics data for each body parts of *T. crocea^a^***

^a^ Lipid classes were grouped in color. For lipid class nomenclature see, http://prime.psc.riken.jp/compms/msdial/lipidnomenclature.html

^b^ DGTS and DGTA are indistinguishable with the analytical method employed here as they are structural isomers to each other [7].

^c^ Galactosyl lipids and free fatty acids (FA) were not quantified, and thus marked ‘-‘

**Table S2. LC-MS ion intensities for plant derived metabolites**

**Table S3. Semi-quantitative lipidomics data:** Grand average of each lipid species was ordered to show 15 most abundant DGCC species in each organ of *T. crocea*.

**Table S4. Semi-quantitative lipidomics data. Grand average of each lipid species was ordered to show 15 most abundant PC species in each organ of *T. crocea*.**

**Table S5. NMR data for GCC and polar portion of DGCC obtained in the present study along with those of DGCC reported** [6].

