## Supplemental data for "Smart utilization of betaine lipids in giant clam *Tridacna crocea*"

### Supplementary Scheme

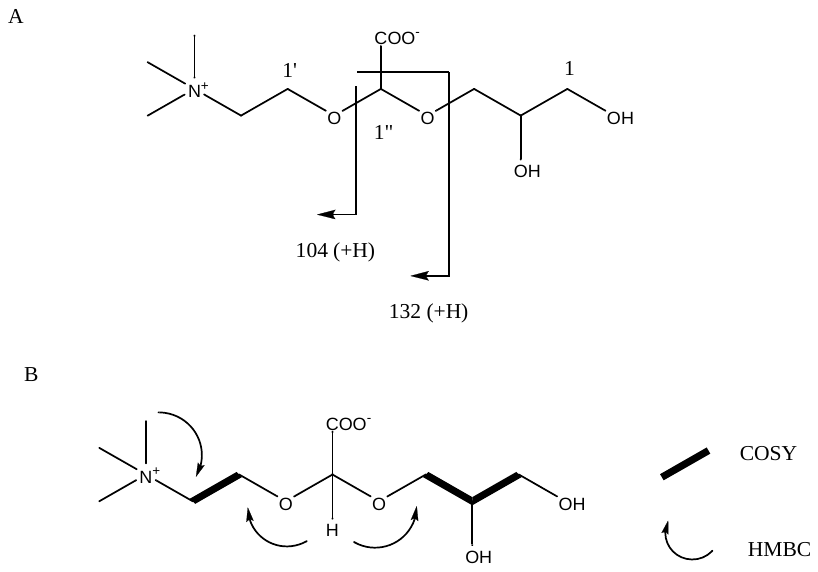

#### Scheme S1. Planer structure of GCC.

(A) fragmentations in ESIMS/MS. (B) Correlation NMR data for GCC.

### Supplementary Figures

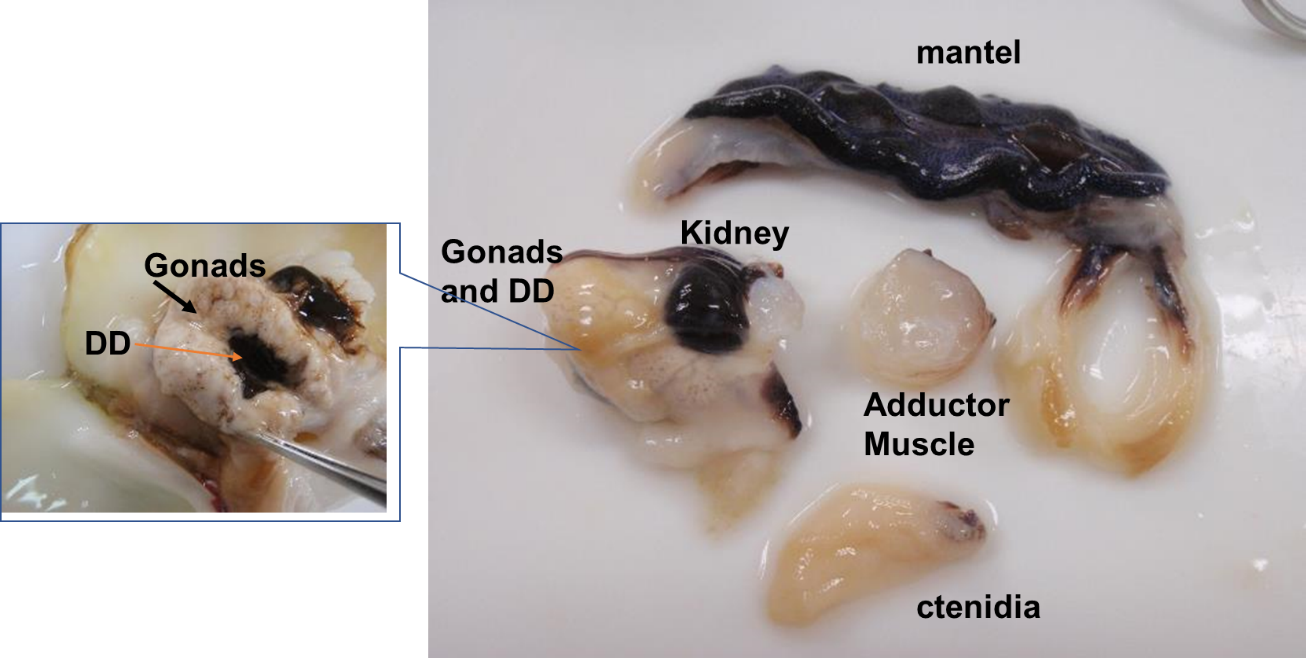

#### Fig. S1. Dissected clam (*Tridacna crocea)* specimen, indicating all body parts used for LC-MS analyses.

Adductor muscle is covered by a thin reddish-brown membrane containing Symbiodiniaceae cells. To avoid contamination of algal cells, we removed this membrane from adductor muscle specimens.

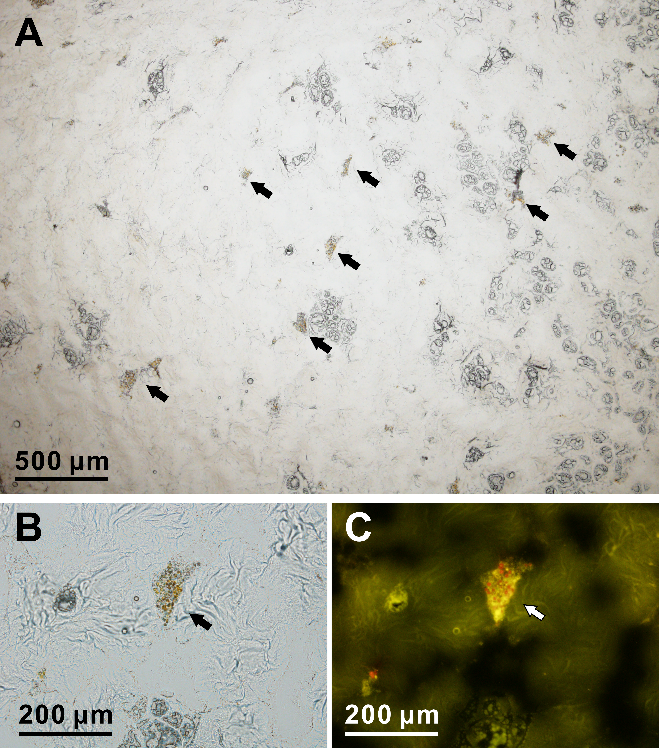

#### Fig. S2. Light micrographs for gonad region.

Small brown patches (arrows) were observed in the gonad region. B, C high magnification observations. Under blue right excitation, a few algal cells were recognized within the patches (C).

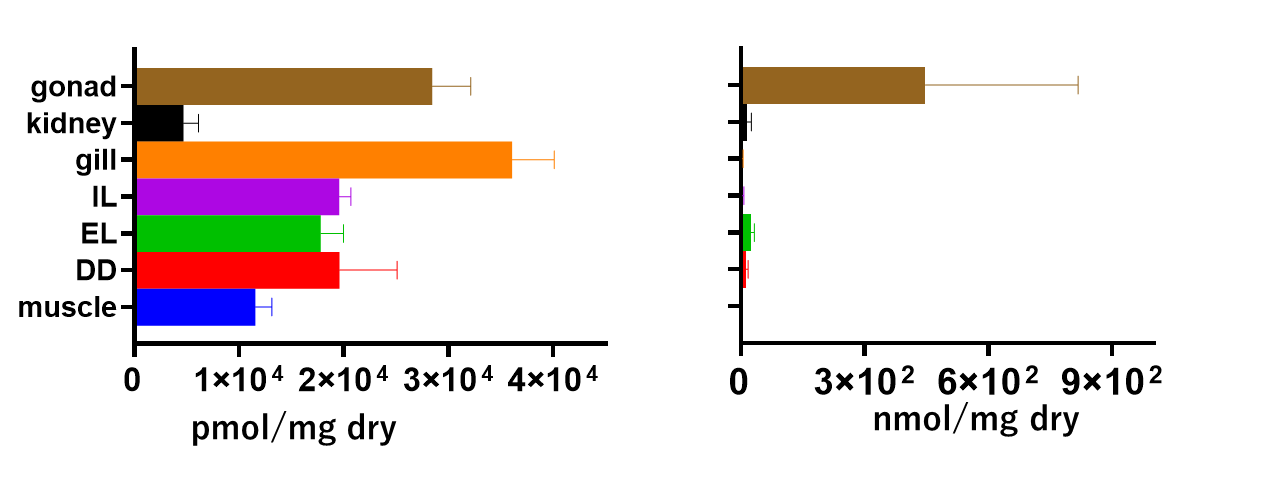

#### Fig. S3. Amounts ether-PE (left) and storage lipids TG (right).

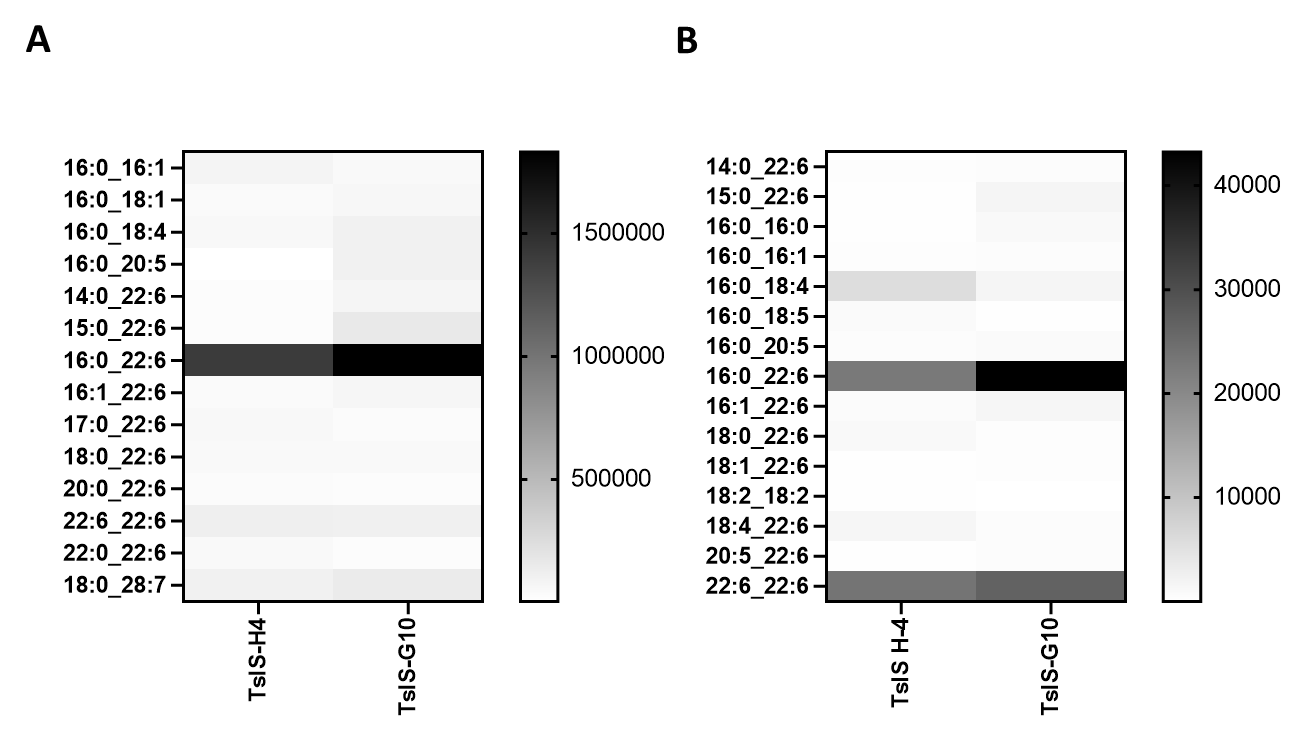

Fig. S4. Heatmap for DGCC (left) and PC (right) in Symbiodiniaceae culture strains (TsIS-H4, and TsIS-G10). Positive ion mode was used for DGCC analysis while negative mode was employed for PC data acquisition because substantially fewer peaks were obtained in the opposite polarity in both classes.

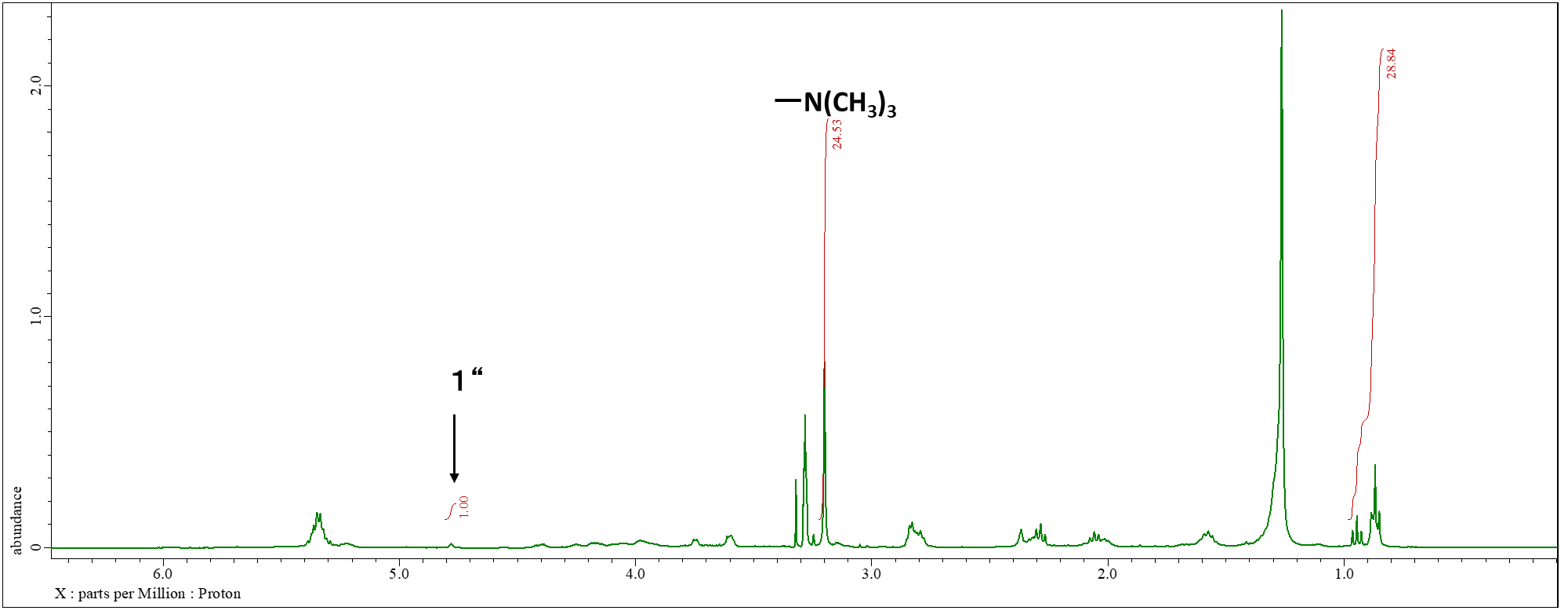

#### Fig. S5. ^1^H NMR (400 MHz) for the DGCC-containing fraction taken in CD_3_OD.

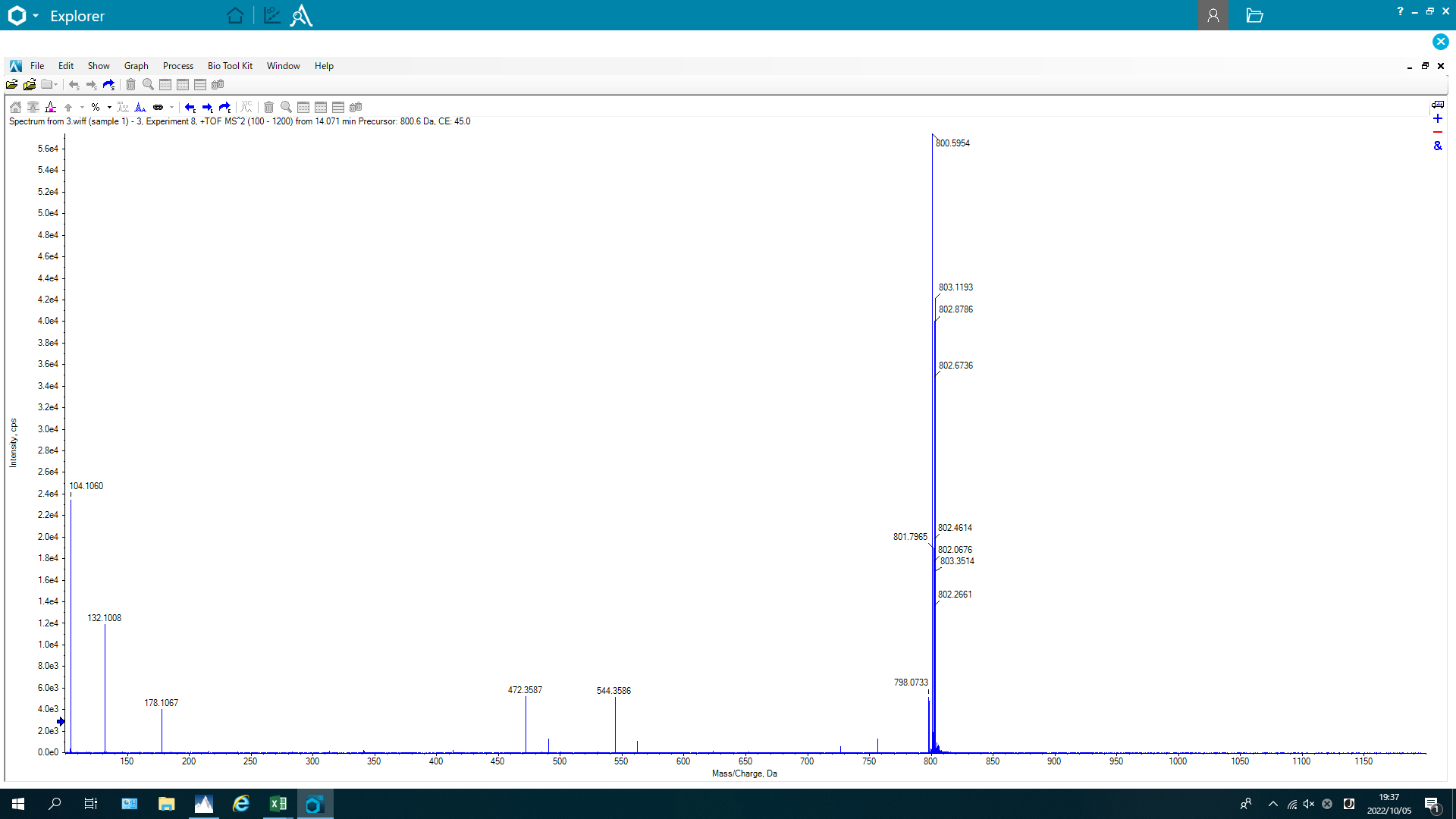

#### Fig. S6. ESIMS for DGCC 22:6_16:0.

A

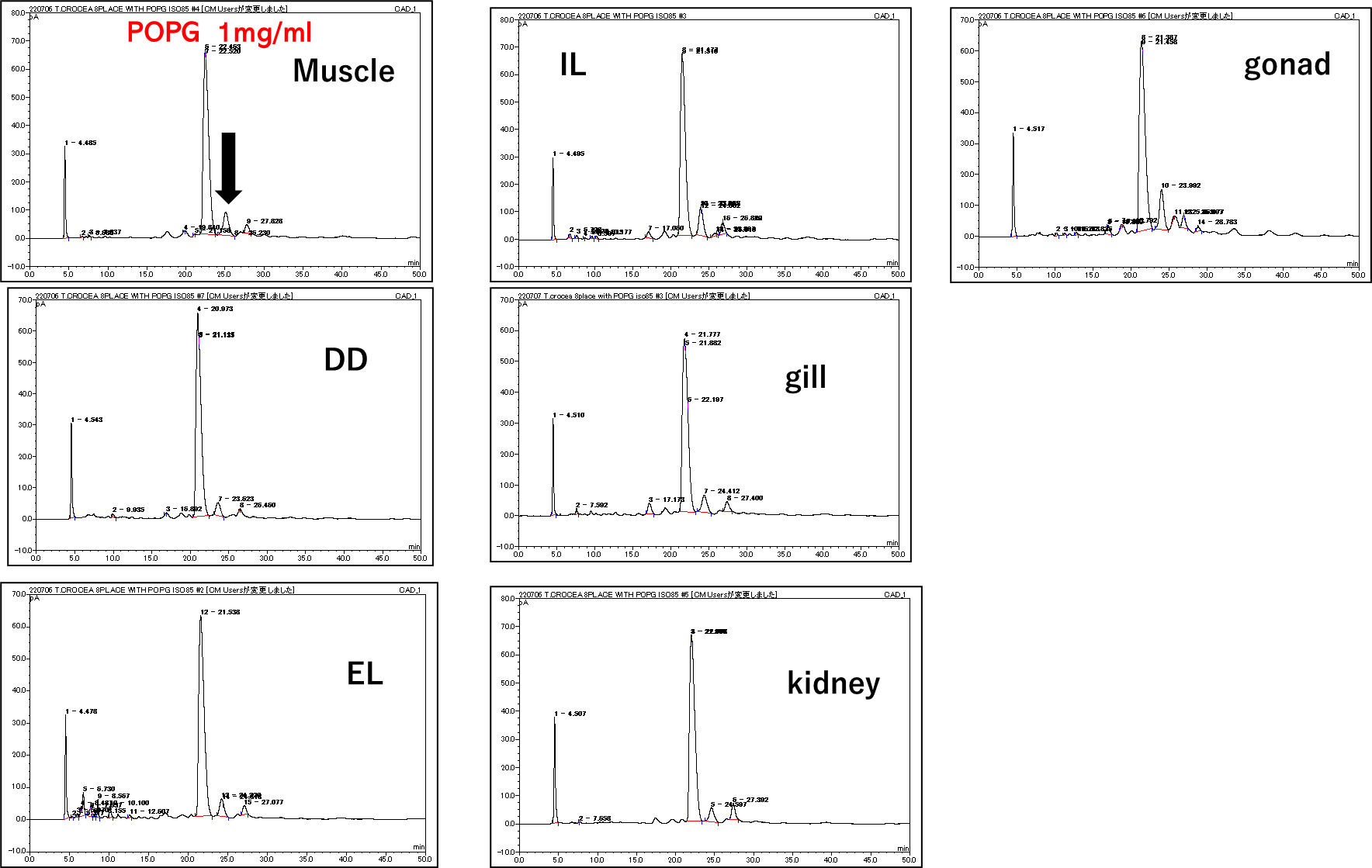

B

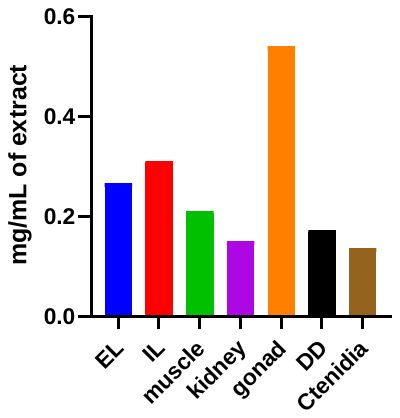

#### Fig. S7. CAD analysis of DGCC in each body part of *T. crocea*.

(A) HPLC-CAD trace of each extract. A large peak at *T*_R_ around 22 min is POPG (1 mg/mL), and DGCC eluted right after the standard (black arrow). The area of concentration of DGCC was calculated from the area of standard and DGCC. One sample was employed to obtain this data due to sample availability.

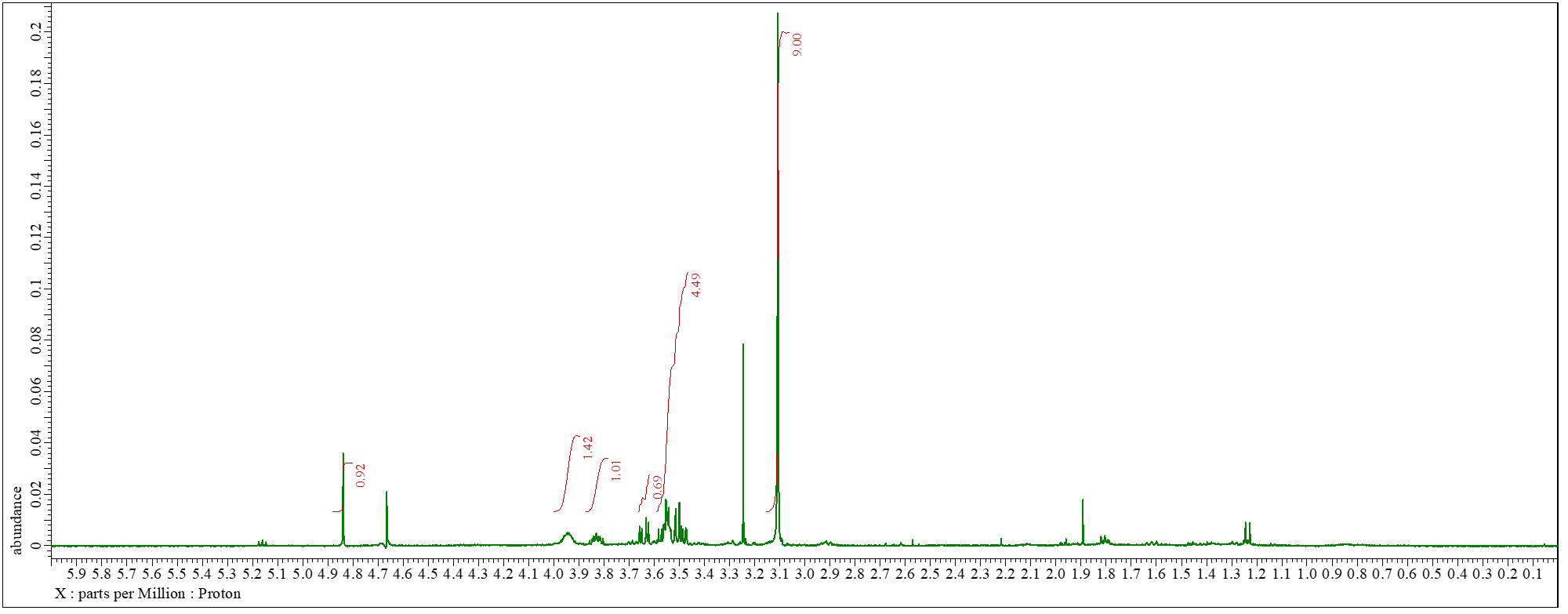

#### Fig. S8. ^1^H NMR (400 MHz) of GCC.

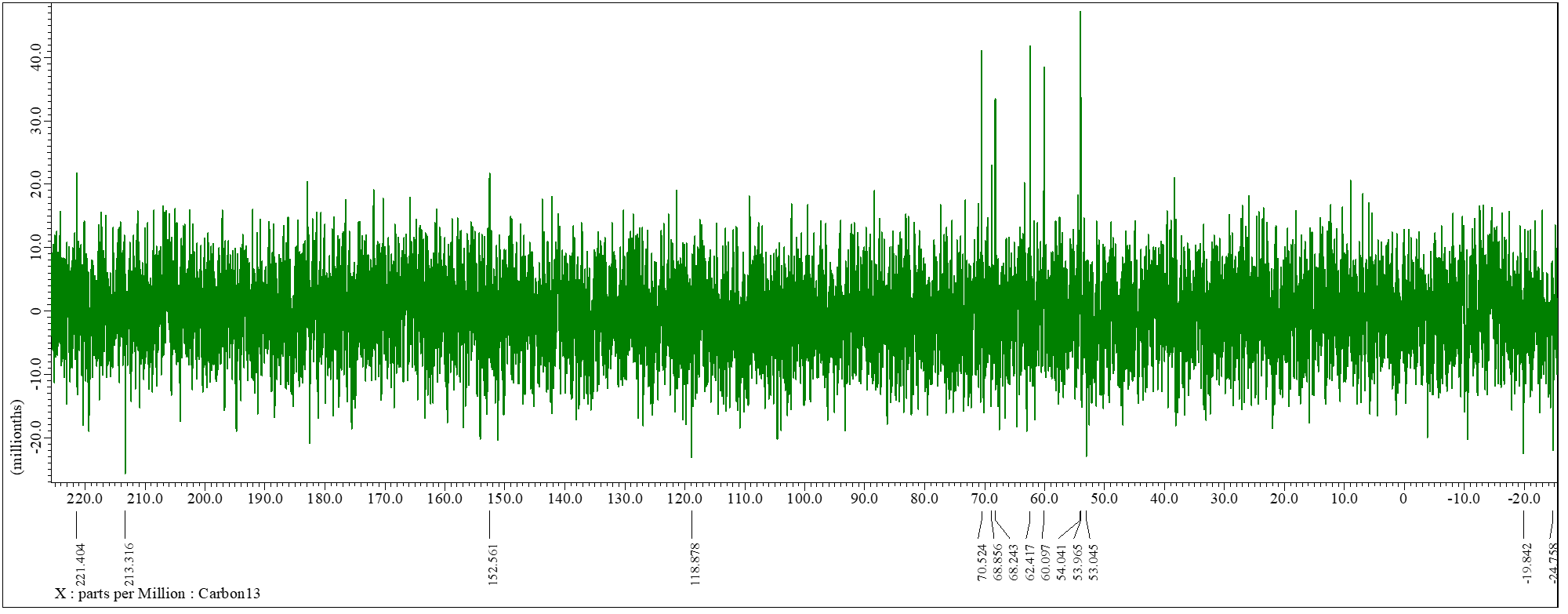

#### Fig. S9. ^13^C NMR (100 MHz) of GCC in D_2_O.

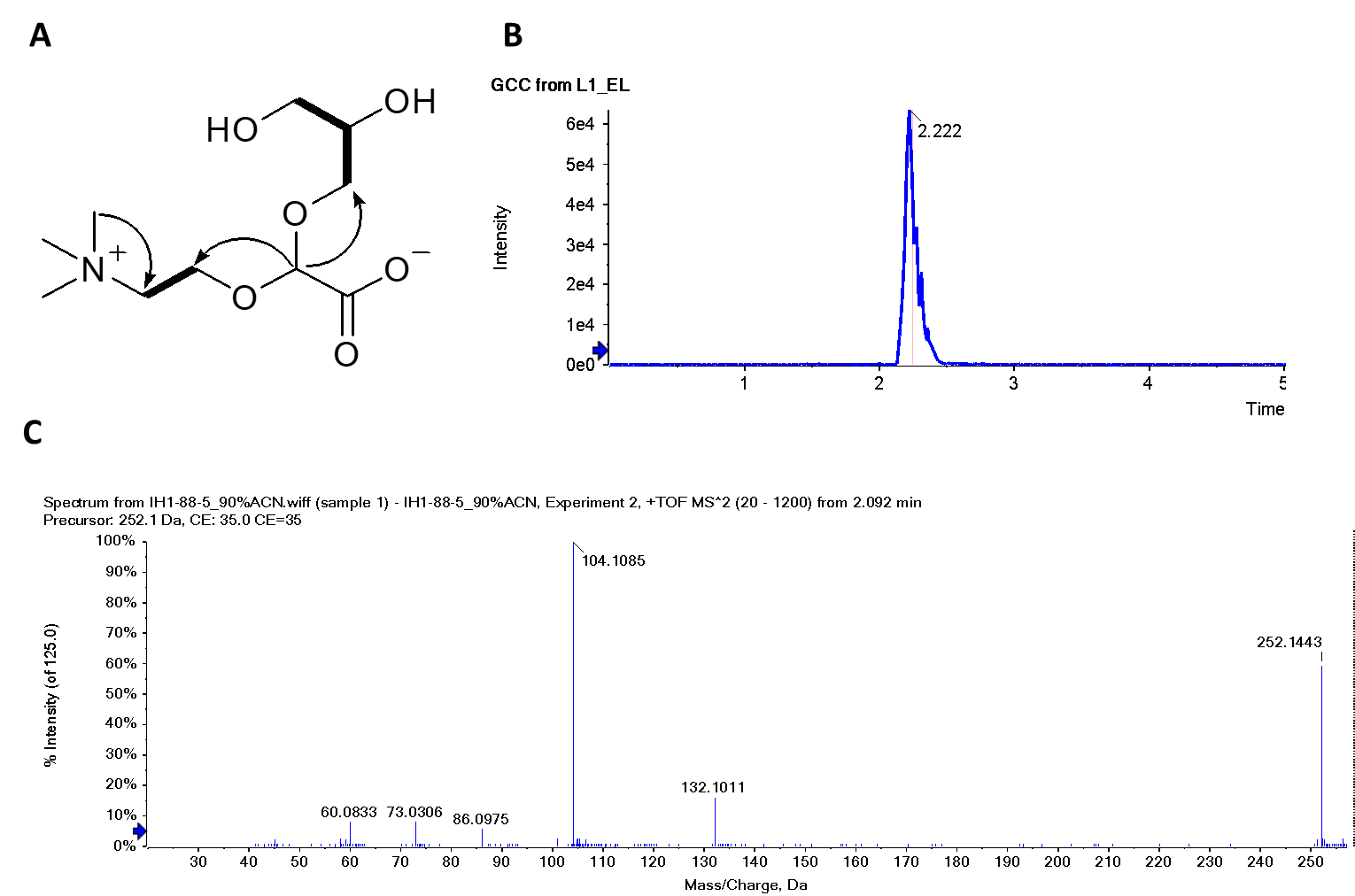

#### Fig. S10. ESIMS/MS data for GCC.

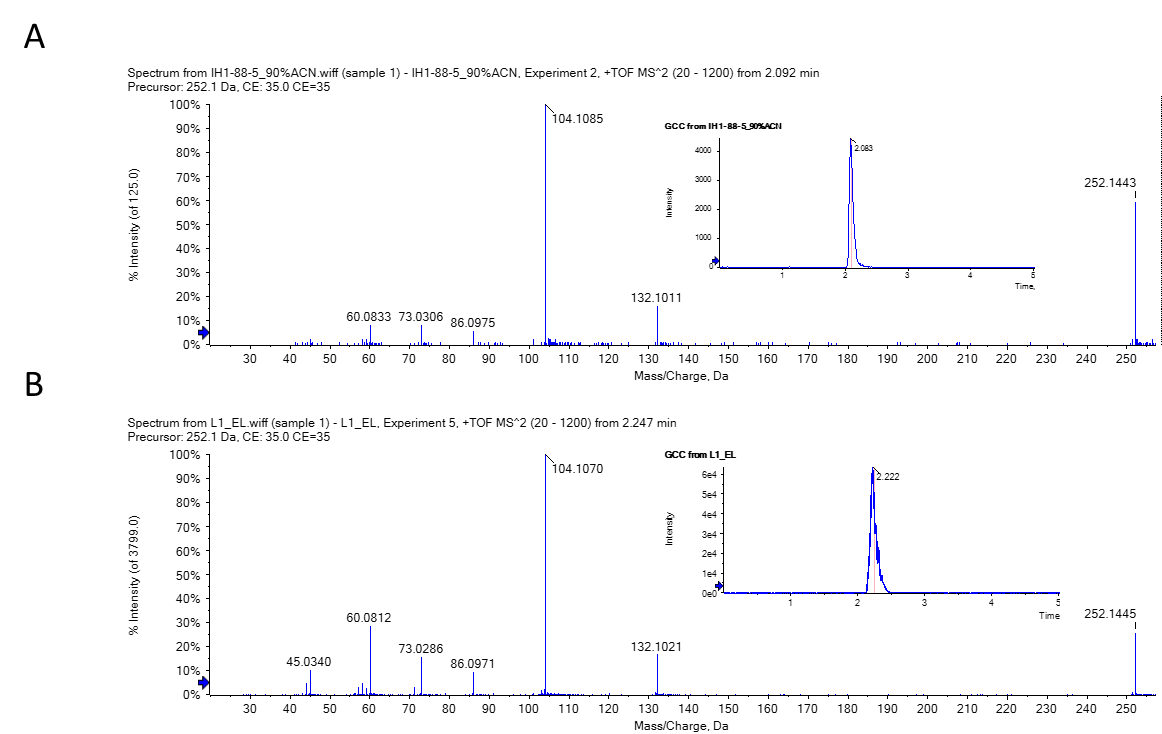

#### Fig.S11 LC-MS analyses of GCC from *Symbiodinium* (A) and *T. crocea* (B).

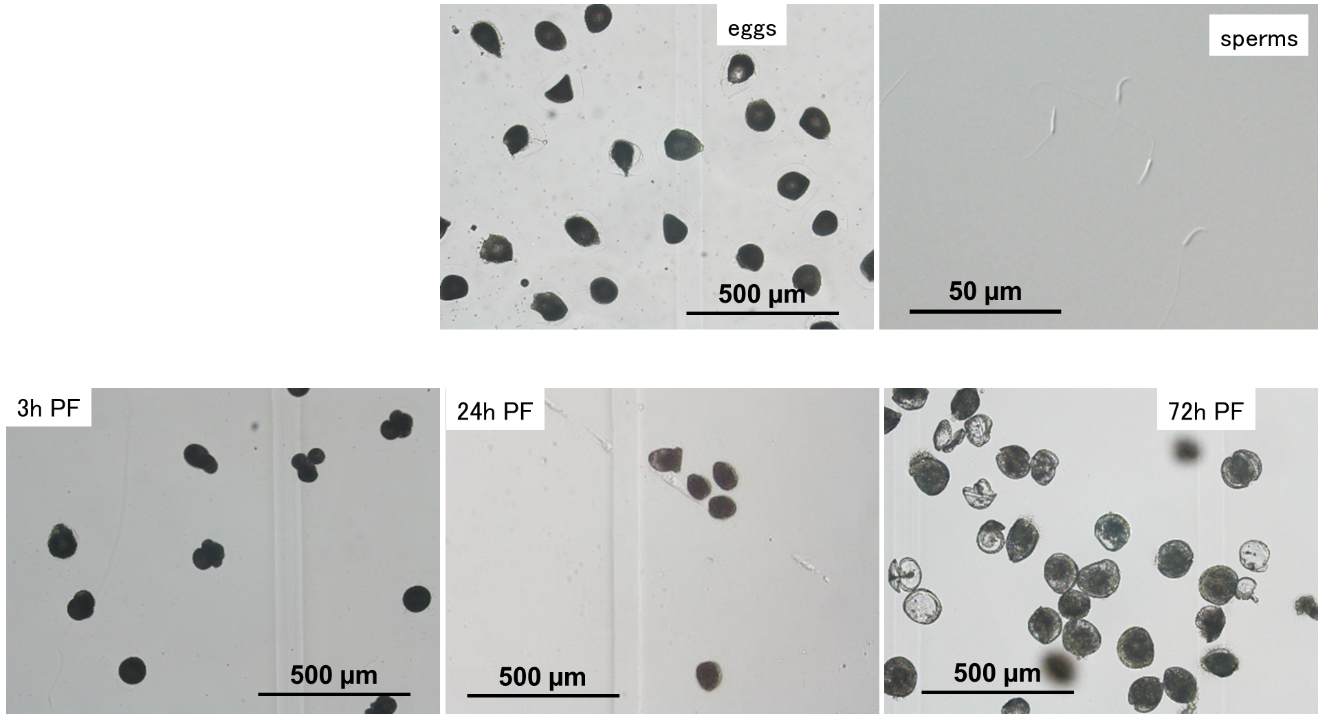

#### Fig. S12. Eggs, sperms, and larvae of *Tridacna crocea* in each developmental stage.

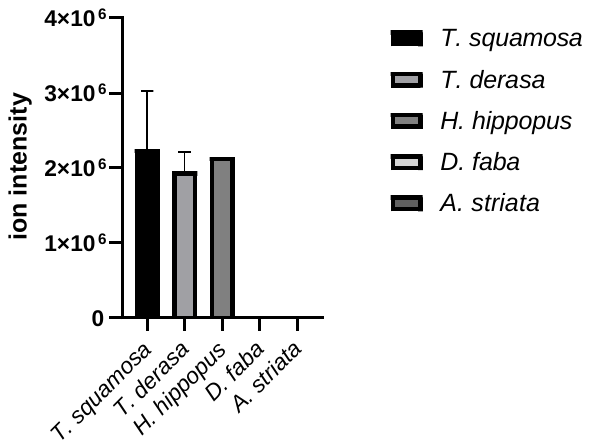

#### Fig. S13. Ion intensities for DGCC in giant clams and other bivalves.

*T. squamosa* (n = 3), *T. derasa* (n = 3), *Hippopus hippopus* (n = 1), and bivalves *Donax faba* (n = 3), and *Atactodea striata* (n = 3).

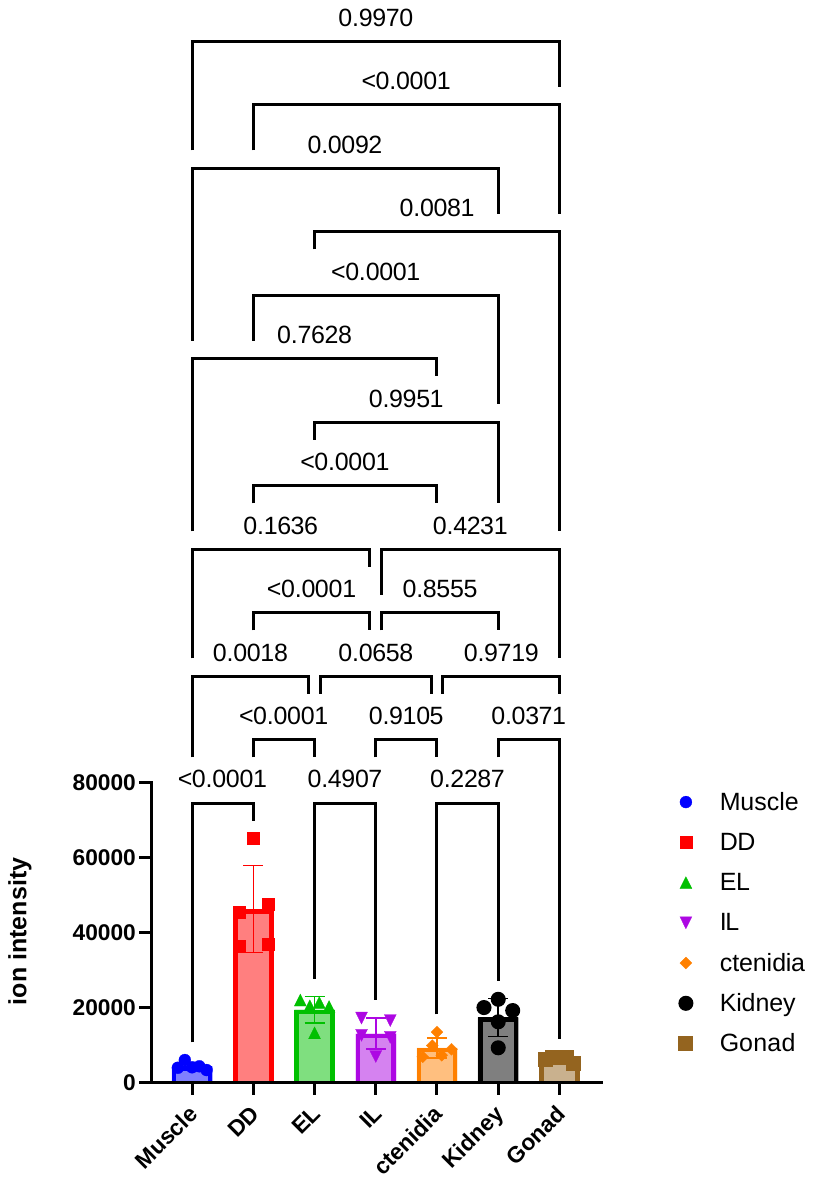

#### Figure S14. Ion intensity for GCC in each organ.

N = 5, One-way ANOVA followed by a Dunnett’s test for multiple group comparison. *P* values are indicated.

### Supplementary Tables

#### Table S1. A summary of semi-quantitative lipidomics data for each body parts of *T. crocea^a^*

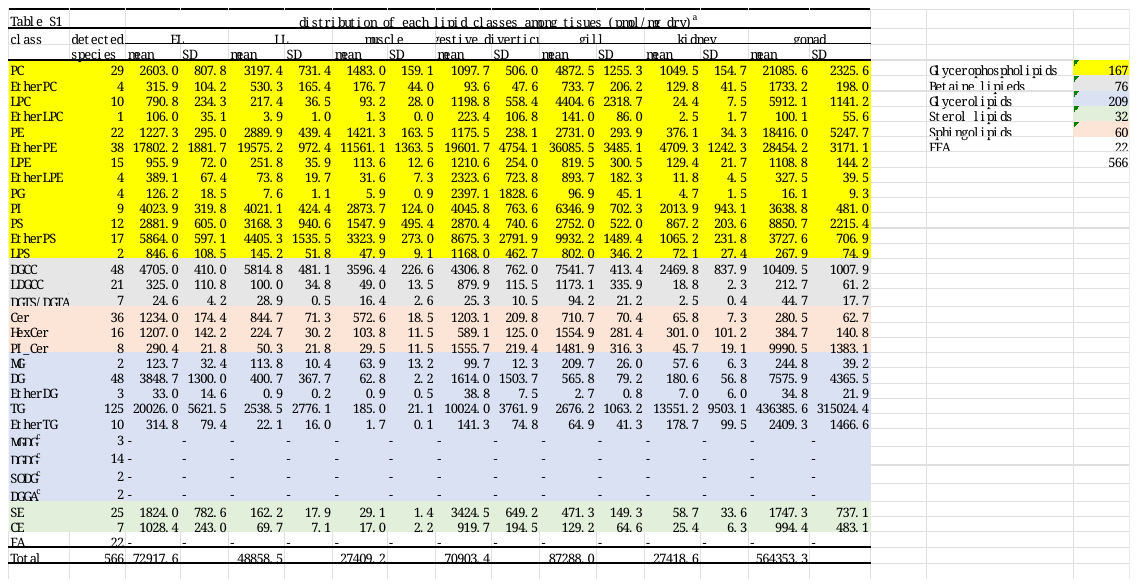

^a^ Lipid classes were grouped in color. For lipid class nomenclature see, <http://prime.psc.riken.jp/compms/msdial/lipidnomenclature.html>

^b^ DGTS and DGTA are indistinguishable with the analytical method employed here as they are structural isomers to each other [6].

^c^ Galactosyl lipids and free fatty acids (FA) were not quantified, and thus marked ‘-‘

#### Table S2. LC-MS ion intensities for plant derived metabolites

|  | muscle | DD | EL | IL | ctenidia | kidney | gonads |
| --- | --- | --- | --- | --- | --- | --- | --- |
| MGDG | 4 | 1033 | 2331 | 155 | 145 | 31 | 49 |
| DGDG | 28 | 11040 | 91252 | 1807 | 687 | 49 | 84 |
| SQDG | 30 | 8485 | 11840 | 719 | 321 | 38 | 222 |
| Peridinin | 11 | 6973 | 12837 | 615 | 154 | 8 | 28 |

DD, digestive diverticula; DGDG,digalactosyldiacylglycerol; EL, epidermal layer; IL, inner layer; LC-MS, liquid chromatography-tandem mass spectrometry; MGDG, monogalactosyldiacylglycerol; SQDG, sulfoquinovosyl diacylglycerols

#### Table S3. Semi-quantitative lipidomics data: Grand average of each lipid species was ordered to show 15 most abundant DGCC species in each organ of *T. crocea*.

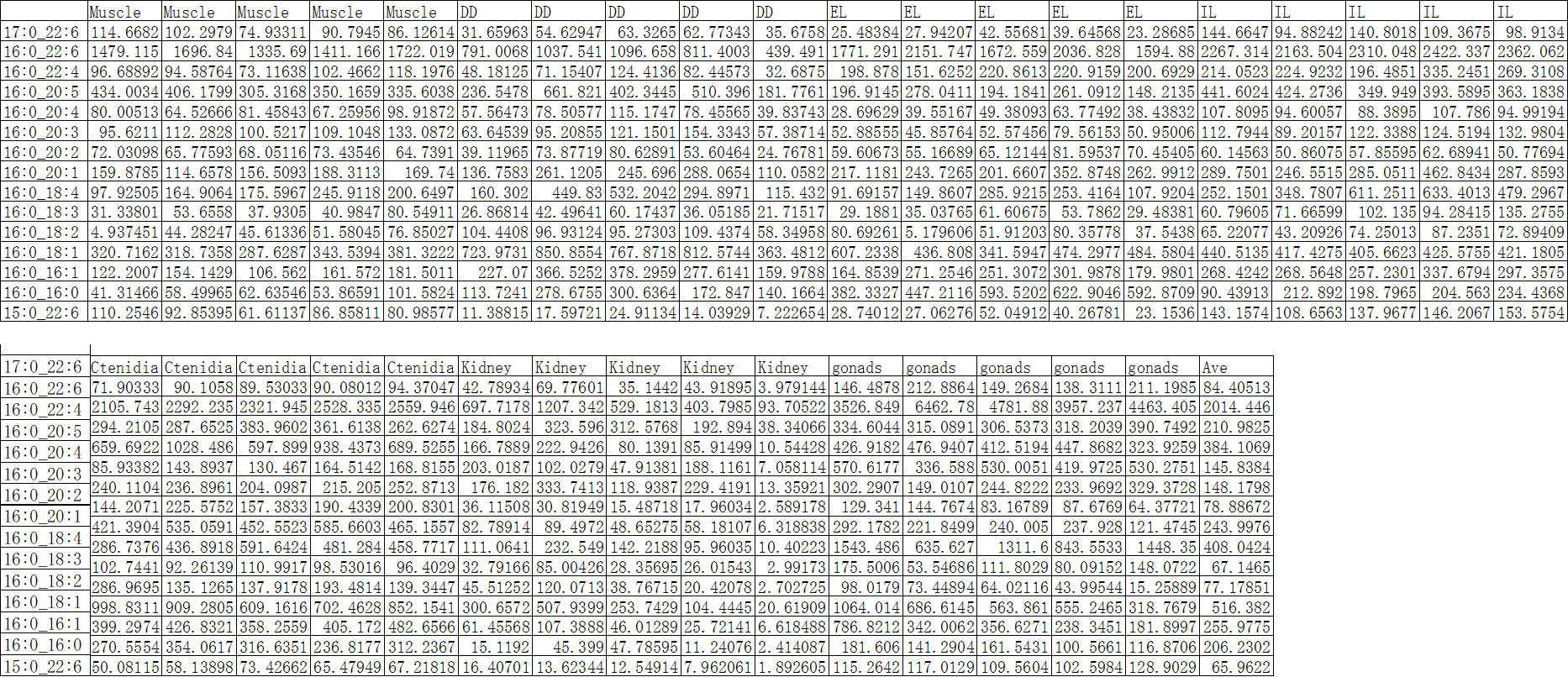

#### Table S4. Semi-quantitative lipidomics data. Grand average of each lipid species was ordered to show 15 most abundant PC species in each organ of *T. crocea*.

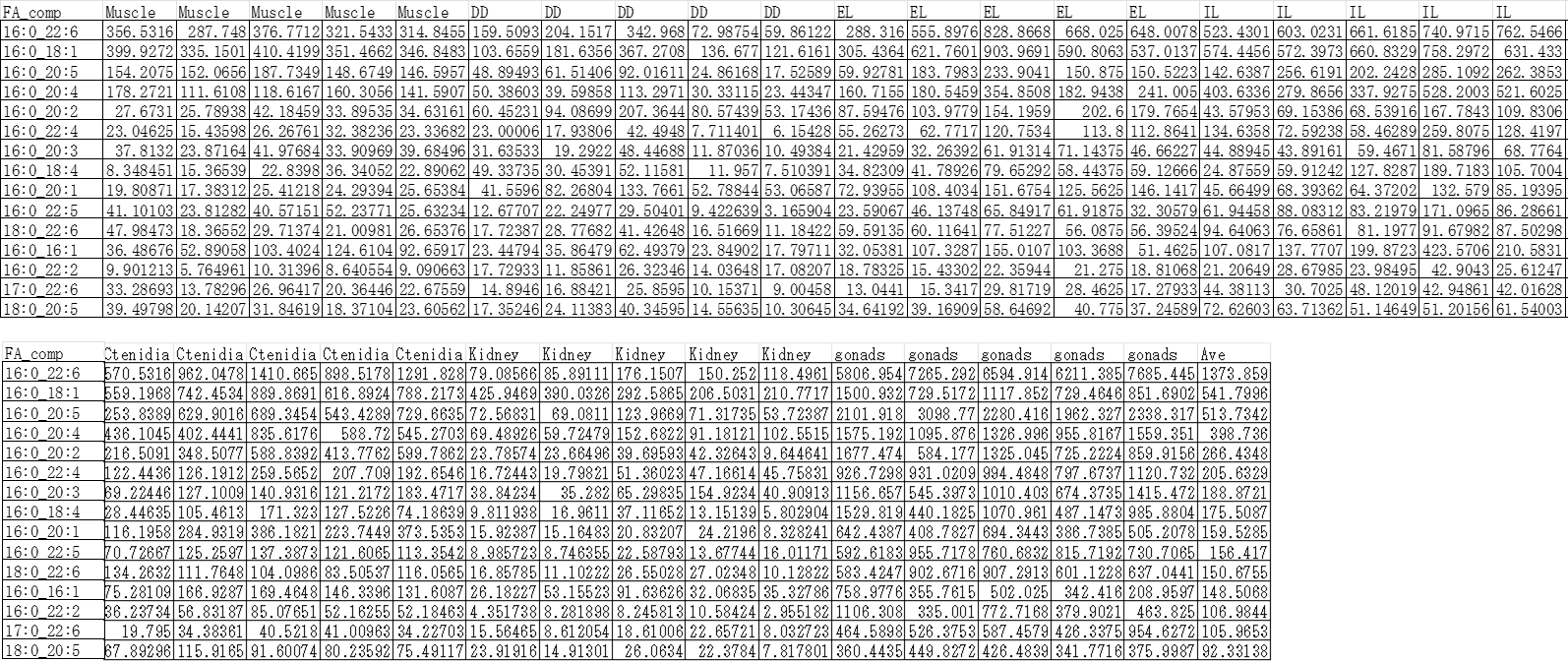

#### Table S5. NMR data for GCC and polar portion of DGCC obtained in the present study along with those of DGCC reported [7].

| Number of  carbon | GCC (D_2_O) | | DGCC (CD_3_OD) | | DGCC [7]  CDCl_3_-CD_3_OD, 1:2 | |
| --- | --- | --- | --- | --- | --- | --- |
| 1 | 3.55, m, 2H | 62.4 | 4.23, 4.38 |  | 4.21, 4.46 | 63.6 |
| 2 | 3.83 | 70.3 | 5.2 |  | 5.27 | 71.4 |
| 3 | 3.64, 3.51 | 68.3 | 3.73 |  | 3.76 | 66.3 |
| 1' | 3.53, m, 2H | 65.5 | 3.57 |  | 3.6 | 66.7 |
| 2' | 3.95 | 60.1 | 3.95, 4.03 |  | 3.93, 4.08 | 60.3 |
| NCH_3_ | 3.11 s, 9H | 54.1 | 3.2 | 52.4 | 3.23, s, 9H | 54.8 |
| 1" | 4.85s, m, | 99.6 | 4.78 |  | 4.78 | 101.4 |
| 2" |  | - |  |  |  | 172,4 |
